# Using OPMs to measure neural activity in standing, mobile participants

**DOI:** 10.1101/2021.05.26.445793

**Authors:** Robert A. Seymour, Nicholas Alexander, Stephanie Mellor, George C. O’Neill, Tim M. Tierney, Gareth R. Barnes, Eleanor A. Maguire

**Affiliations:** Wellcome Centre for Human Neuroimaging, UCL Queen Square Institute of Neurology, University College London, London WC1N 3AR, UK

**Keywords:** Auditory evoked field, OP-MEG, wearable neuroimaging, beamformer, motion capture

## Abstract

Optically pumped magnetometer-based magnetoencephalography (OP-MEG) can be used to measure neuromagnetic fields while participants move in a magnetically shielded room. Head movements in previous OP-MEG studies have been up to 20 cm translation and ∼30° rotation in a sitting position. While this represents a step-change over stationary MEG systems, naturalistic head movement is likely to exceed these limits, particularly when participants are standing up. In this proof-of-concept study, we sought to push the movement limits of OP-MEG even further. Using a 90 channel (45-sensor) whole-head OP-MEG system and concurrent motion capture, we recorded auditory evoked fields while participants were: (i) sitting still, (ii) standing up and still, and (iii) standing up and making large natural head movements continuously throughout the recording – maximum translation 120 cm, maximum rotation 198°. Following pre-processing, movement artefacts were substantially reduced but not eliminated. However, upon utilisation of a beamformer, the M100 event-related field localised to primary auditory regions. Furthermore, the event-related fields from auditory cortex were remarkably consistent across the three conditions. These results suggest that a wide range of movement is possible with current OP-MEG systems. This in turn underscores the exciting potential of OP-MEG for recording neural activity during naturalistic paradigms that involve movement (e.g. navigation), and for scanning populations who are difficult to study with stationary MEG (e.g. young children).

## 1. Introduction

Magnetoencephalography (MEG) is a non-invasive neuroimaging technique that measures small magnetic fields outside of the head originating from current flows throughout the brain (Cohen, 1968). MEG data have very high temporal resolution, allowing the characterisation of evoked responses and neuronal oscillations at the sub-millisecond timescale (Baillet, 2017). Unlike the electrical potentials measured with electroencephalography, magnetic fields are largely unaffected by signal distortions from the conductivity profiles of scalp tissue. Until recently, the only sensors routinely used for MEG were superconducting quantum interference devices (SQUIDs). However, these sensors require cryogenic cooling using liquid helium, rendering MEG systems stationary and expensive.

A new generation of wearable MEG sensors called optically pumped magnetometers (OPMs) have been developed (Boto et al., 2018), that measure small magnetic fields (see Tierney et al., 2019 for a review) and have a similar sensitivity to SQUID systems (7-15 ft/Hz from 1-100 Hz) but, crucially, do not require cryogenic cooling. This means that the sensors can be placed closer to the scalp, resulting in up to five-fold signal magnitude increases over conventional SQUID systems (Boto et al., 2016; Iivanainen et al., 2017). Another advantage of OPMs over SQUIDs is that sensors can be placed independently on the scalp surface rather than held in rigid arrays. The exact location of the sensors can, therefore, be customised on a participant-by-participant basis, rather than being fixed for one specific head shape and size. Furthermore, because sensors can move with the head, data can be collected during participant movement, and with minimal constraints on recording time, which will significantly improve the utility of MEG both as a clinical tool (Vivekananda et al., 2020) and for basic neuroscientific research (e.g. Barry et al., 2019; Boto et al., 2021; Roberts et al., 2019). This is because cohorts who find the head immobilisation associated with SQUID-MEG and magnetic resonance imaging (MRI) challenging (e.g. children) can be scanned more easily, and naturalistic paradigms that involve participant movement can be more readily deployed.

OPMs have been used to successfully measure a variety of neuromagnetic fields while participants were seated, including: beta-band (13-30 Hz) oscillations in motor cortex during movement (Boto et al., 2018), theta-band (4-8 Hz) oscillations from the hippocampus while the participant was unconstrained (Barry et al., 2019; Tierney et al., 2021a), and visual gamma (40-80 Hz) oscillations (Iivanainen et al., 2020), auditory evoked fields (AEFs) (Borna et al., 2017), somatosensory evoked fields (Boto et al., 2017) and median nerve evoked fields (Iivanainen et al., 2019) while participants were stationary. Further miniaturisation of OPMs has facilitated the construction of more lightweight, whole-head sensor arrays (Hill et al., 2020), capable of measuring resting-state connectivity comparable to that of a 275-channel SQUID system while using 50 OPM sensors (Boto et al., 2021).

Despite the promise of OPM-based MEG (OP-MEG), recording neuromagnetic fields during participant movement comes with substantial challenges. In a typical magnetically shielded room (MSR) that comprises multiple layers of mu-metal, there remains a static field component (due to both the room itself and the external static field) of the order of nano-Tesla. Any small sensor translations or rotations relative to this background magnetic field within the MSR will cause large signal change in the recorded data. Without correction, the magnitude of these artefacts easily exceeds any neural signals of interest and can even exceed the dynamic range of current OPM sensors (±5.56 nT for QZFM second generation sensors; this limit is applied by the manufacturer, QuSpin, to avoid gain errors due to non-linearity in the OPM response as the magnetic field increases - see Tierney et al., 2019), resulting in periods of unusable data.

Recent work has focussed on minimising background fields through a variety of complementary techniques. First, on-board coils allow the cancellation of static fields inside an OPM sensor’s cell (Osborne et al., 2018); this works well for stationary OPM arrays. Second, all of the initial OPM measures showing robustness to participant motion used external biplanar coils and reference magnetometers to dynamically null the background fields around a participant’s head to just ∼0.5 nT (Boto et al., 2018; Hill et al., 2020; Holmes et al., 2018, 2019). This technique works well and is essential for OPMs to function in the majority of MSRs that were originally designed for conventional SQUID-MEG. However, the biplanar coils arrangement reduces the amount of possible effective movement to 40 cm x 40 cm x 40 cm with current designs. Finally, the passive shielding requirements of OPM systems, unlike SQUID systems, extend down to 0 Hz. More recent MSRs designed specifically for OPMs and equipped with degaussing coils (Altarev et al., 2015) can reduce these static magnetic fields to just ∼1.5 nT at the room centre (Hill et al., 2020). It is also worth noting that despite the progress made in reducing static background fields within MSR environments, movement artefacts in modern OP-MEG systems can still be similar in magnitude to the neuromagnetic fields of interest (Boto et al., 2018; Holmes et al., 2018). Fortunately, these artefacts typically have spatiotemporal properties distinct from neuromagnetic fields. Signal-processing techniques based on spatial and temporal filtering, such as beamforming (Brookes et al., 2008; Hillebrand & Barnes, 2005; Van Veen et al., 1997), can therefore be used for interference suppression during participant movement (Tierney et al., 2021b).

Neuroscientific work has demonstrated the ability of OP-MEG to detect neuromagnetic fields at the same time as naturalistic head movement while seated. For example, Boto et al. (2018) showed beta-band power modulations during a finger abduction paradigm while making natural head movements, including drinking tea, accompanied by a maximum head displacement of ±10 cm. Holmes et al. (2018) detected lateralised visual evoked fields during head translation of ±9.7 cm and rotation of ±34°. Similar visual evoked results were reported using a virtual reality set-up (Roberts et al., 2019). In addition, the first paediatric OP-MEG study showed beta-band motor responses associated with head movements of 4.73 ± 1.21 cm translation and 28.1° ± 6.71° rotation (Hill et al., 2019). These results represent a step-change over stationary SQUID-MEG systems, in which movements over 0.5 cm can severely degrade data quality (Gross et al., 2013). While movement compensation algorithms (e.g. based on signal space separation, Medvedovsky et al., 2007) are available for head movements on the order of 0.5-6 cm during SQUID-MEG scanning, much larger naturalistic movements (over ∼6cm) are not possible with a stationary array of MEG sensors.

In the current study, we sought to push the movement limits of OP-MEG even further to determine whether it is technically possible to measure neuromagnetic fields while participants are standing up and deliberately making very large continuous rotations and translations of the head. Collecting OP-MEG data from participants standing up compared to sitting down is technically challenging because the sensors are typically located away from the centre of the room (where shielding is optimal), where remnant field gradients are much steeper, thereby exacerbating movement related artefacts. If OP-MEG is feasible in situations where participants are standing up and making continuous head movements, it would open up opportunities for a range of naturalistic MEG paradigms. This could include studies involving interactions with objects or touch-screens (Jungnickel & Gramann, 2016), or using omnidirectional treadmills to simulate exploration of virtual environments (Schiza et al., 2019).

As a proof-of-principle, the experimental paradigm was simple for this study: auditory tones were used to elicit AEFs. These arise in supratemporal auditory cortex, and typically comprise a small 50 ms response (M50), and larger 100 ms (M100) and 200 ms (M200) responses (for a review see Hari, 1990). Due to their reliability, AEFs are often used in the context of MEG for benchmarking purposes (e.g. Sekihara et al., 2002; Taulu & Hari, 2009; Tierney et al., 2021b). AEFs are also a low-frequency phenomenon (1-40 Hz), that overlap in the frequency domain with movement artefacts (<10 Hz). Therefore, in this study, the measurement of AEFs while participants were standing and moving represented an ideal test case for spatiotemporal interference suppression techniques.

OP-MEG data were acquired while head motion was tracked using a six-camera motion capture system. We show that with appropriate pre-processing and artefact rejection steps, and in combination with beamforming, it is possible to measure AEFs while participants are standing and making very large, continuous translations (maximum = 120 cm) and rotations (maximum = 198°) across the course of an experiment including, on average, ∼5 cm of movement during individual 0.5 second trials.

## 2. Methods

### 2.1 Participants

Two healthy, right-handed males, aged 26 and 29, participated in the study. They both provided written informed consent and the study was approved by the University College London Research Ethics Committee.

### 2.2 OP-MEG data acquisition

OP-MEG data were acquired in an MSR (Magnetic Shields Ltd) located at University College London. The room has internal dimensions of 438 cm x 338 cm x 218 cm and is constructed from two inner layers of 1 mm mu-metal, a 6 mm copper layer, and then two external layers of 1.5 mm mu-metal.

A total of 45 dual-axis OPMs (QuSpin Inc., QZFM second generation) were used in the study, which have a sensitivity of ∼15 fT/√Hz between 1-100 Hz. Forty three sensors (i.e. 86 channels) were placed in a participant-specific 3D-printed “scanner-cast” (Boto et al., 2017), which was designed using the participant’s structural MRI scan (Chalk Studios), see Fig. 1A. The scanner-cast was designed to keep the sensors in slots fixed in relation to the brain during participant movement, and to minimise co-registration errors (Meyer et al., 2017). As each scanner-cast was printed by first creating a 3D image in the same coordinate space as participant’s structural MRI brain scan, a sensor’s position and orientation could be calculated offline relative to the slot in which it was placed. Sensor position was set as the centre of the cell of the OPM sensor, which was slightly offset from the physical centre. As shown in Fig. 1A, custom plastic clips were used to arrange the OPM sensor ribbon cables for effective cable management. In addition, the larger cables were organised into bundles and fixed to a wearable backpack, to facilitate participant comfort during movement. The sensors were arranged to evenly cover the whole head (Fig. 1B), with an approximately symmetrical layout across each hemisphere of the brain. An additional two sensors were placed away from the participant to provide reference signals, however they were not used for interference suppression in this study.

**Fig. 1.**
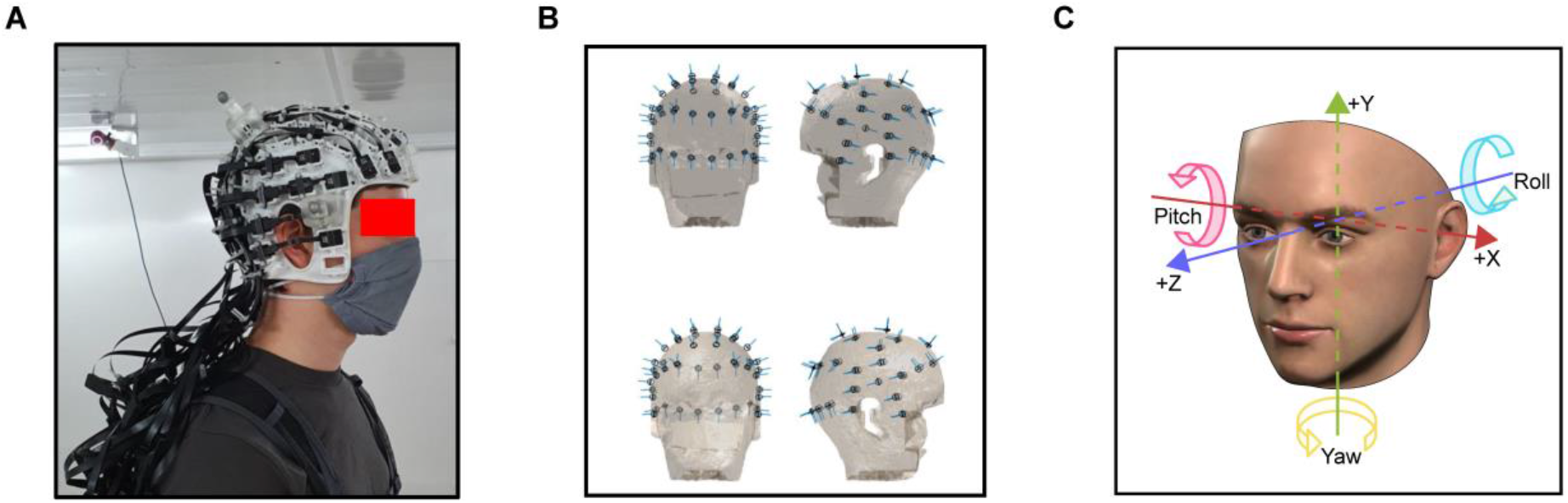
**(A)** The experimental setup. Inside the magnetically shielded room OPM sensors were secured in a 3D printed scanner-cast with cables fixed into a backpack. **(B)** The location of the sensors for participant 1, plotted on a 3D mesh derived from the participant’s de-faced structural MRI scan. Sensitive axes are shown by the blue lines. **(C)** The six degrees of freedom used to describe the translation and rotation of the rigid body.

Before the start of the experiment, the MSR was degaussed to minimise the residual magnetic field in the MSR. Before the start of each experimental run, the OPM sensors were calibrated using a manufacturer-specific procedure. This involves energising coils within an OPM to produce a known field, and the output of the sensor is then measured and calibrated to this known field. These calibration values, determined at the start of the experiment, may become suboptimal as sensors move during the course of the experiment. However we did not find any evidence that this impacted the quality of our data in this study.

Data were recorded using a 16-bit precision analog-to-digital converter (National Instruments) with a sample rate of 6000 Hz. For our setup, the two sensitive OPM axes were oriented both radial and tangential to the head, and had the effect of increasing the dimensionality of the data and facilitating spatial interference suppression methods (Brookes et al., 2021; Tierney et al., 2021b). This resulted in 90 channels of OPM data.

### 2.3 Paradigm

An auditory evoked response paradigm, adapted from Garrido et al. (2008), was used. The auditory tones had a duration of 70 ms (5 ms rise and fall times) and frequency of 500-800 Hz in steps of 50 Hz. An auditory tone of the same frequency was presented 2-8 times before randomly switching to an auditory tone of a different frequency. We collapsed our analysis across auditory tones of all frequencies. The inter-stimulus interval was 0.5 s (no stimulus jitter was included). Stimuli were presented via PsychoPy (Peirce, 2009), through MEG-compatible ear tubes with Etymotic transducers, and the volume was adjusted to a comfortable level as specified by the participant.

In the first run of data acquisition, the participant was seated on a plastic chair in the centre of the MSR and was instructed to keep as still as possible. In the second run, the participant stood upright in the middle of the MSR, again keeping as still as possible. This run was included because the gradient in the background magnetic fields is steeper when a participant stands up and positions their head away from the centre-point (at ∼1.35 m height) of the MSR. In the third run, the participant was instructed to stand in the middle of the MSR and continually move and rotate their head in any direction they wished. In each run, ∼570 individual auditory tones were presented, resulting in 275-300 s worth of OP-MEG data.

### 2.4 Head position tracking

For head position tracking, an array of six OptiTrack Flex13 (NaturalPoint, Inc.) motion capture cameras were used. These cameras were placed in three corners of the room, with two in each (one high up, one low down), to allow for complete coverage of the tracking area. Six retro-reflective markers were attached to the scanner-cast in multiple fixed positions to form a “rigid body”. These were tracked passively using the OptiTrack cameras at 120 Hz throughout the experiment. By measuring the joint translation of markers on the rigid body, the motion capture system could calculate the position and orientation of the rigid body while the participant moved within the MSR. Note that the physical occlusion of one or multiple markers due to participant movement produced gaps in the data where no information about the position of a marker was known. These gaps were interpolated to produce uninterrupted motion capture data using two methods. First, for any gap, if three or more of the six markers on the rigid body were visible, this permitted “pattern based” interpolation to determine the only possible position of the occluded marker(s). After pattern based interpolation, some gaps remained - in runs 1 and 2, the still conditions (sitting and standing), 0.86 % of the data remained as gaps (maximum duration 1.2 s; mean duration 26 ms). In run 3, the moving condition, there were more gaps, 8.94 %, but of similar duration (1.24 s maximum; 29 ms mean). Remaining gaps, up to 0.83 s in duration (100 samples at 120 Hz), were interpolated using a second method, cubic spline fit to the data either side of the gap. Pattern based interpolation was then applied again, removing all remaining gaps. To remove high frequency noise caused by marker vibrations, the trajectory data were low-pass filtered at 2 Hz using a 4^th^ order Butterworth filter applied bidirectionally to achieve zero-phase shift. The rigid body was then solved based on the continuous trajectory data of the six markers. Both the marker trajectory data and the solved rigid body data were visually inspected for errors that can occur during processing (e.g. mislabelling of markers, jumps in the data) and corrections were made where necessary. Finally, the motion capture position and orientation data were up-sampled and synchronised with the OP-MEG data.

### 2.5 Data analysis

All analyses were performed in MATLAB 2019a using the Fieldtrip Toolbox (Oostenveld et al., 2011), SPM (Litvak et al., 2011) and custom scripts, which can be found openly online at: https://github.com/FIL-OPMEG/movement_auditory_ERF.

### 2.6 OP-MEG pre-processing

Data were down-sampled to 1000 Hz for computational efficiency. A multiple linear regression was performed to reduce the magnetic field artefacts covarying with head position tracking data (Holmes et al., 2018). The regression included the rigid body position (X, Y, Z) and rotation data (pitch, yaw, roll; Fig. 1C), and was performed in overlapping 10-second-long windows.

Next, the data were cleaned using homogenous-field correction which approximates magnetic interference as a spatially constant field, on a sample-by-sample basis (see Tierney et al., 2021b for further details). This is calculated from the sensors’ orientation information, rather than position, and is removed from the data via linear regression. To avoid ringing artefacts from notch filters (de Cheveigné & Nelken, 2019), we instead used a spectral interpolation procedure (Leske & Dalal, 2019) to remove high amplitude peaks in the frequency spectrum corresponding to: (i) 50 Hz power-line contamination and its harmonics, and (ii) 120 Hz and 83 Hz contamination from the motion capture camera system. Specifically, the 120 Hz originates from the infrared LEDs on the OptiTrack cameras operating at this frequency, and the 83 Hz peak is a result of temporal aliasing when the 120 Hz interference is digitally sampled alongside the 923 Hz OPM modulation signal. The OPM data were then high-pass filtered at 2 Hz using a 5^th^ order Butterworth filter applied bidirectionally to achieve zero-phase shift.

We next performed manual visual artefact detection in 5 s chunks to remove any remaining sections of data still contaminated by: (i) field changes greater than 100 pT (over the course of the 5 s chunk) from participant movement and environmental interference not successfully removed by the preceding steps, (ii) intermittent high frequency noise, which may have been due to the OPM cables rubbing against each other, and (iii) high amplitude ‘steps’ in the data present across all channels. For participant 1: 3.1% of trials were lost from run 1, 0.6% of trials from run 2, and 4.7% of trials from run 3. For participant 2: 1.4% of trials were lost from run 1, 5.3% of trials from run 2, and 7.6% of trials from run 3. Finally, data were low-pass filtered at 40 Hz using a 6^th^ order Butterworth filter applied bidirectionally, and then epoched into trials of 0.7 s (−0.2 s pre-stimulus; 0.5 s post-stimulus onset). Low-pass filtering was performed after visual data inspection so that high-frequency artefacts could be more easily identified. To ensure equitable power across runs (sitting, standing, standing and moving), comparisons were made using 525 of the remaining good trials for each run (randomly selected).

For all runs, the results of the initial pre-processing steps are shown in Supplementary Fig. S1. For runs 1-2 (sitting and standing), the combination of homogenous-field correction and high-pass filtering both reduced the maximum field change from 29.2 pT to 3.4 pT (sitting) and 48.2 pT to 4.5 pT (standing). However, the movement data regression step increased the maximum field change present in each trial. This is potentially due to the movement data containing noise which is introduced into the data via regression when little is actually present. For this reason, we re-ran the pre-processing for runs 1-2, but not run 3, avoiding the movement data regression step. These data were used to generate Figs. 4-6.

For the moving condition, see Supplementary Fig. S1 (right), each pre-processing step helped to reduce the low-frequency interference in the data, including movement data regression. The maximum field change in each trial was reduced from an average of 654.7 pT to 242.1 pT following movement regression, to 87.8 pT following homogenous-field correction (Tierney et al., 2021b), and to 9.8 pT following the high-pass filter.

### 2.7 Sensor-level analysis

Data were averaged and baseline corrected using the 0.1 s of data before stimulus onset. Event-related activity was plotted for each sensor, and sensor-level fieldmaps were produced for the evoked magnetic waveforms oriented radially to the head (the tangential components being more difficult to visualise).

### 2.8 Source-level analysis

A participant’s T1-weighted structural MRI scan was used to create a forward model based on a single-shell description of the inner surface of the skull (Nolte, 2003). Using SPM12, a nonlinear spatial normalisation procedure was used to construct a volumetric grid (5 mm resolution) registered to the canonical Montreal Neurological Institute brain.

Source analysis was conducted using a linearly constrained minimum variance (LCMV) beamformer (Van Veen et al., 1997), which applies a spatial filter to the MEG data at each point of the 5 mm grid. Based on recommendations for optimising MEG beamforming (Brookes et al., 2008), for the main analyses a regularisation parameter of lambda = 0.1% was used. Beamformer weights were calculated by combining lead-field information with a sensor-level covariance matrix, computed from the unaveraged single-trial data from 0-0.5 s post-stimulus onset. To counter the bias towards the centre of the head, whole-brain power maps were weighted by the beamformer-projected noise – the neural activity index (NAI) (Van Veen et al., 1997). Due to the highly correlated, near simultaneous neural activity in bilateral auditory regions evoked by auditory stimulation, traditional beamformers often yield suboptimal results for auditory data (Brookes et al., 2007; Sekihara et al., 2002; Van Veen et al., 1997). Consequently, we opted to construct a dual source model in which the beamformer scans pairs of symmetric dipoles across the two hemispheres of the brain (Popov et al., 2018).

Next, we defined a region of interest (ROI) in primary auditory (A1) cortex, using a multi-modal parcellation (Glasser et al., 2016). To obtain a time-course of the data within A1, we performed a principal components analysis on the concatenated filters of each grid-point within the ROI, multiplied by the sensor-level covariance matrix, and extracted the first component (Schoffelen et al., 2017). The pre-processed, sensor-level data were multiplied by this spatial filter to obtain an A1 “virtual channel”. No normalisation (e.g. NAI) was performed for the ROI analysis; instead, a one sample student t-test was conducted at each time point across trials to allow for easier comparison with the sensor-level ERF t-values.

## 3. Results

### 3.1 Head-tracking data

The position (or translation) of the rigid body formed of the head, scanner-cast and OPM sensors can be described via three degrees of freedom: right-left (X), down-up (Y), back-forward (Z). The range of movement in each direction over the course of run 3 (where participants were standing and deliberately moving their head) is shown in Table 1 – these values are far larger than the other two runs that involved sitting still and standing still (data reported in Supplementary Tables S1, S2). For each participant, the continuous position data were plotted as 50-bin histograms (Fig. 2, top panels). This demonstrated that for both participants, right-left and back-forward movement was normally distributed, suggesting that participants were making natural head movements centred on their position at the start of the run. For down-up movement, the histograms show that both participants moved primarily downwards, as expected from a starting standing position. For an alternative view of the position data continuous plotted linearly over time, see Supplementary Fig. S2.

**Table 1.** Rigid body data. Range of values across the standing and moving condition (run 3), for each of the six degrees of freedom.

|  | Participant 1 | Participant 2 |
| --- | --- | --- |
| Right-Left | 101.4 cm | 97.2 cm |
| Back-Forward | 119.9 cm | 114.8 cm |
| Down-Up | 57.1 cm | 39.5 cm |
| Pitch | 111.2° | 115.8° |
| Yaw | 198.1° | 108.3° |
| Roll | 187.9° | 175.2° |

**Fig. 2.**
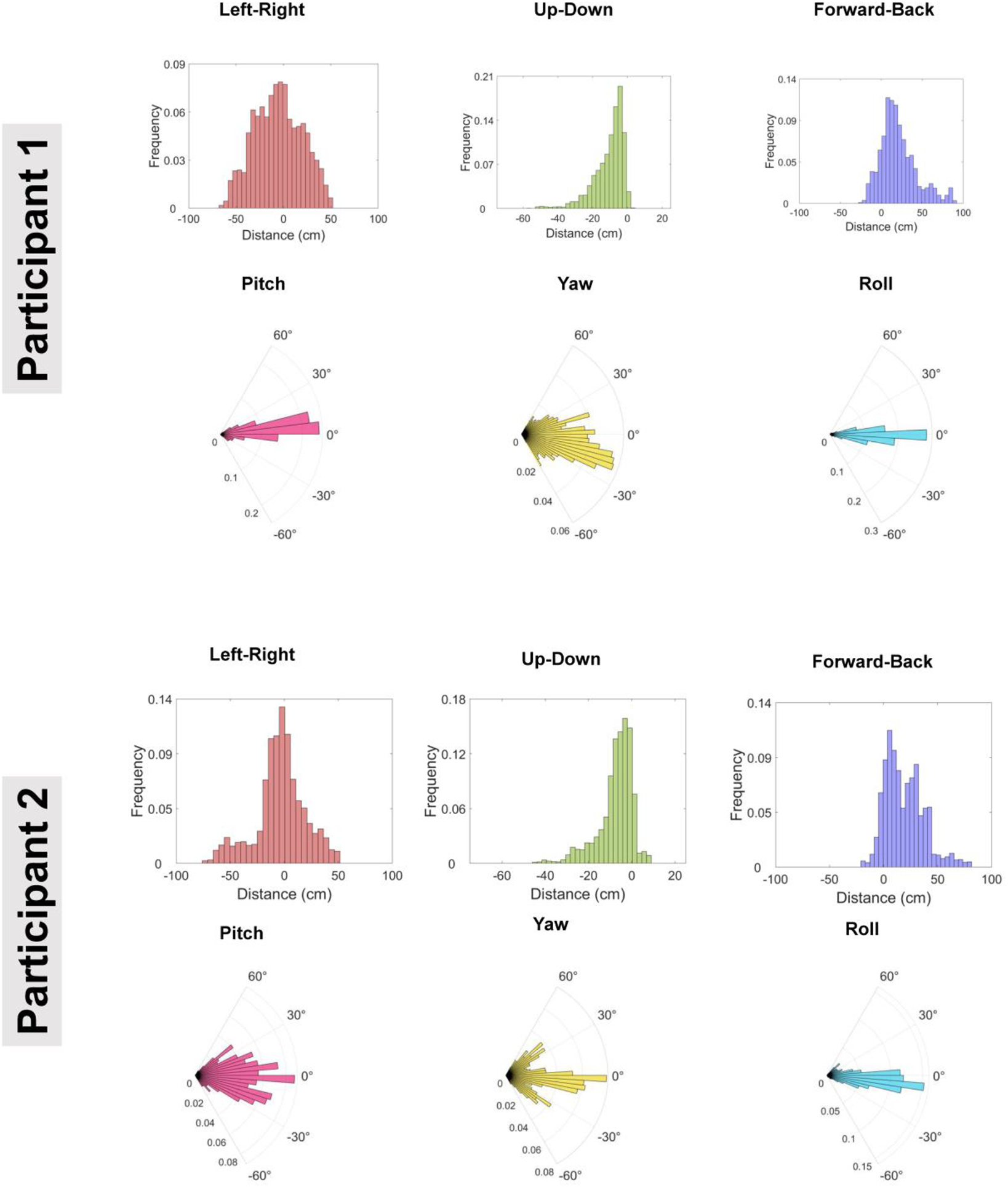
Histograms showing the range of movement across each degree of freedom of the continuous rigid body data. For reference, colours match the transformations shown in Fig. 1C.

The rotation of the rigid body can be described as pitch, yaw and roll (Fig. 1C). The range of rotations over the course of the experiment in each direction are shown in Table 1. These values from the ‘standing and moving’ run were far larger than the other two runs (data reported in Supplementary Tables S1, S2). For each participant, the continuous position data were plotted as polar histograms (Fig. 2, bottom panels for each participant). The vast majority of the rotation data were normally distributed between −60° to +60°. This suggests that both participants made various natural rotations of the head from a forward-facing position. For an alternative view of the rotation data plotted linearly over time, see Supplementary Fig. S2. By combining the translation and rotation data, and applying the transformations to a 3D model, the movement of the rigid body can be animated and visualised in real-time. A 40 s example movie (Supplementary Movie 1) is provided for illustration purposes.

Next, using the motion capture data, Euclidian distance over the course of the experiment was calculated (Supplementary Fig. S3). A bar graph with individual data points was then produced to show the maximum Euclidian distance travelled for each 0.5 s-long trial (Fig. 3). The data clearly show that both participants moved *continuously* during the experiment, with substantial movement on nearly every trial (participant 1 mean distance = 5.1 cm per trial; participant 2 mean distance = 3.9 cm per trial).

**Fig. 3.**
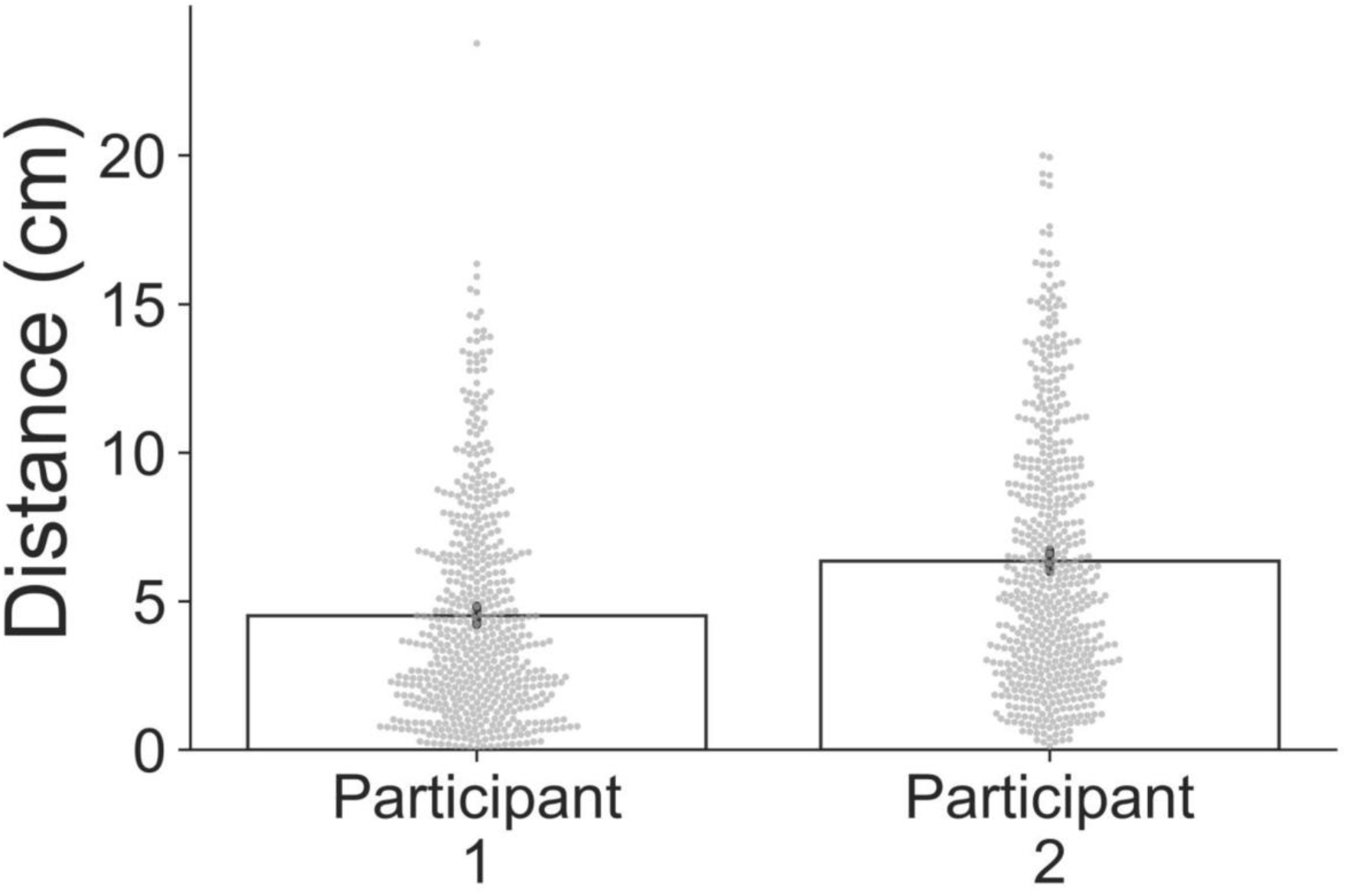
Bar graph showing, for each participant, Euclidian distance of the rigid body travelled during each 0.5 s trial. Error bars correspond to 95% confidence intervals and individual trial data points are shown in grey.

Overall, these motion capture data demonstrate that both participants moved their head continuously throughout the experiment, with large translations/rotations across all six degrees of freedom, mainly centred on their start point.

### 3.2 Sensor-level OP-MEG data

After applying high-pass (2 Hz) and low-pass (40 Hz) filters, and removing trials containing artefacts, the sensor-level data were averaged. Evoked waveforms and fieldmaps are shown in Fig. 4. For both participants, the sitting data and the standing data showed clear evidence of AEFs, including M100 and M200 responses, with bilateral sources evident on the fieldmaps from 0.08-0.12 s. There was some evidence of M100 and M200 responses in the standing and moving data also, however, the data were noisier due to the presence of low-frequency artefacts from participant movement. Focussing on the OPM sensor showing the highest M100 response, the pattern of results is similar - even after pre-processing, AEF t-values were lower when participants were standing and moving their heads, compared with the other two runs (see Supplementary Fig. S4).

**Fig. 4.**
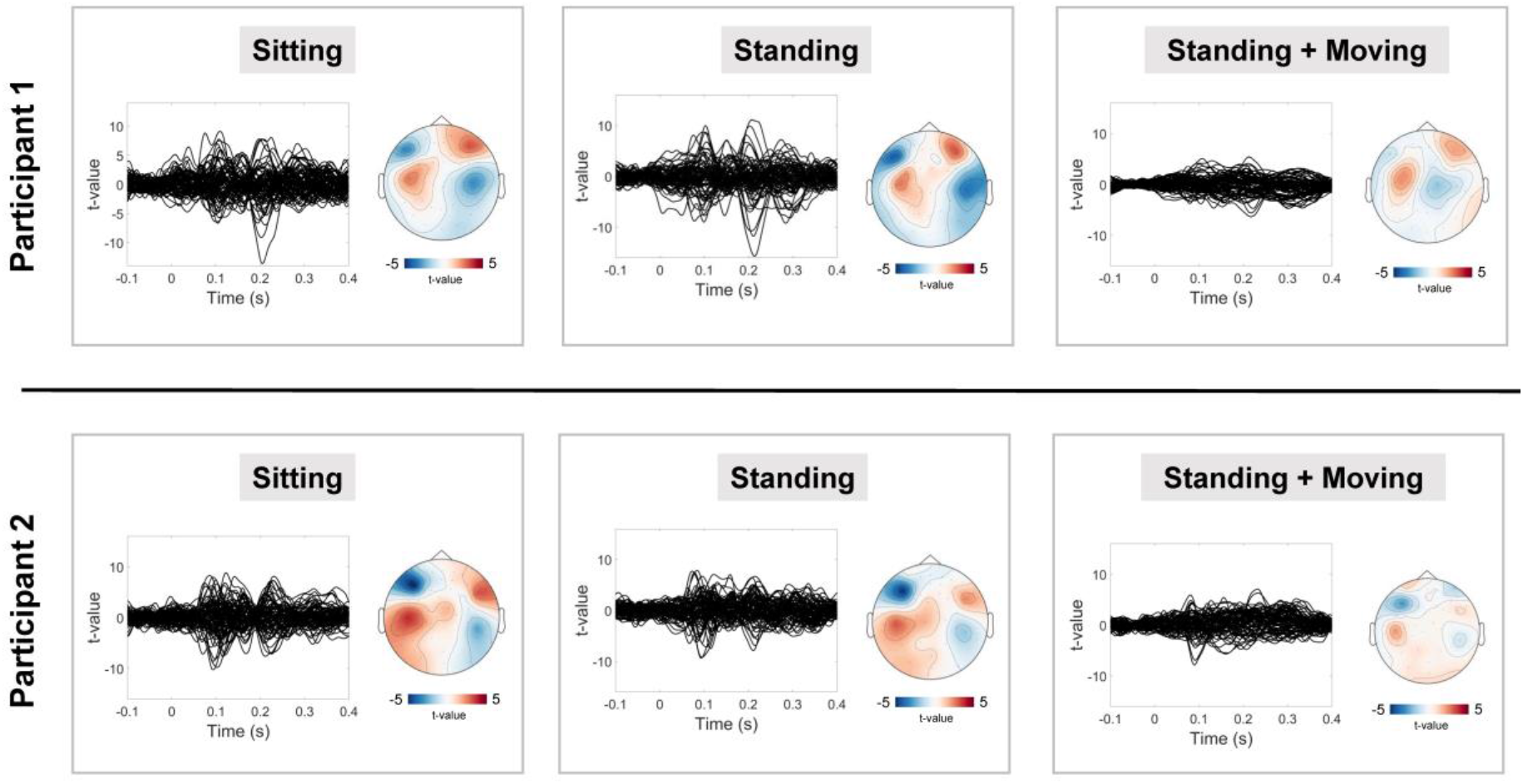
Sensor-level data. For each condition (sitting, standing, standing and moving), evoked waveforms are plotted alongside a 2D fieldmap for data from 0.08 s to 0.12 s post-stimulus onset. The fieldmap only shows magnetic fields oriented radially to the head.

### 3.3 Source-level OP-MEG data

The NAI was averaged for data between 0.08 s to 0.12 s post-stimulus onset, corresponding to the classic M100 AEF (Hari, 1990), and plotted on the SPM canonical brain. Results showed that the evoked magnetic field localised to bilateral auditory cortex, with similar NAI values across all three runs (Fig. 5).

**Fig. 5.**
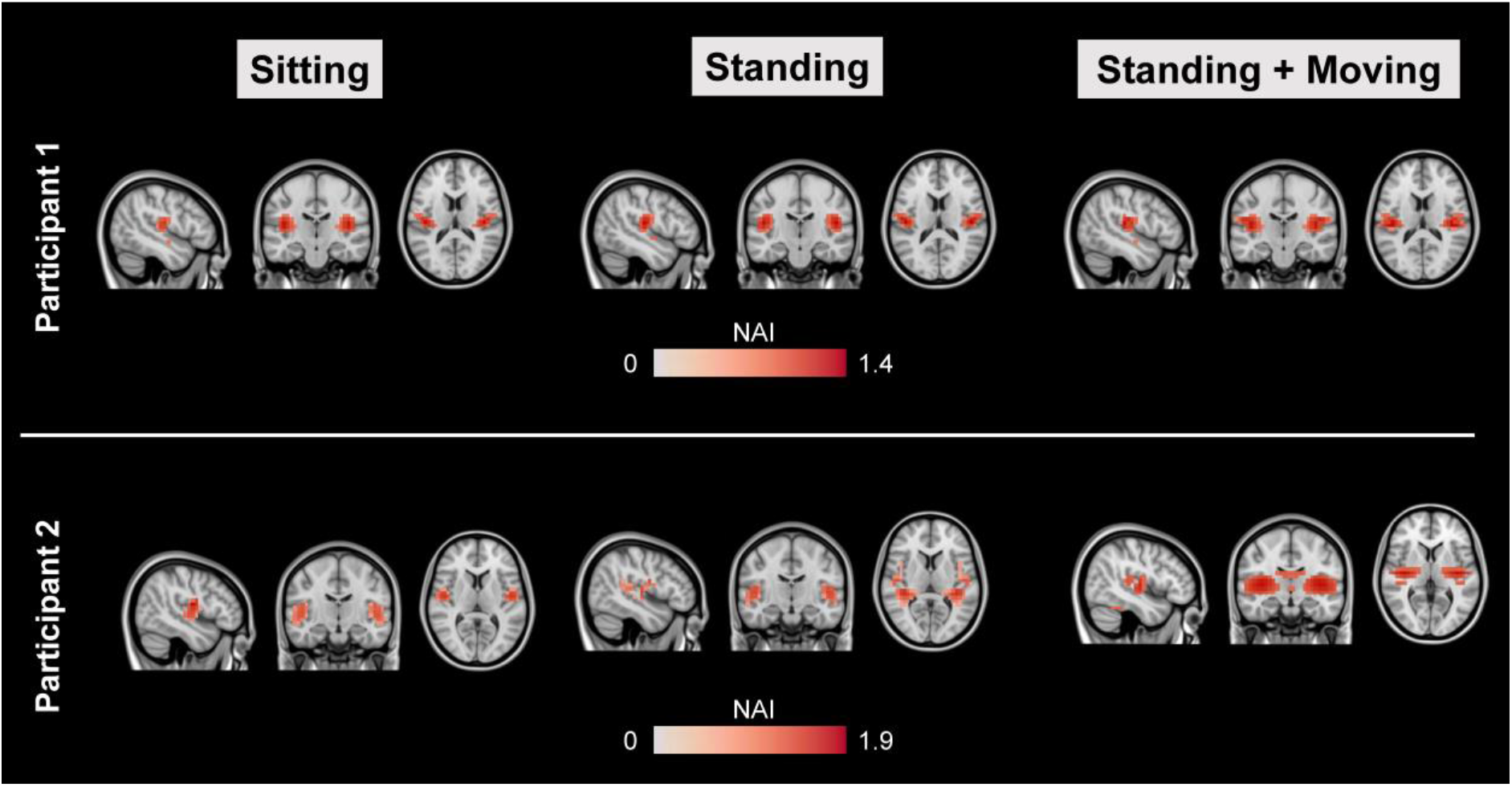
Whole-brain M100 source localisation results. The Neural Activity Index (NAI) was calculated for each run between 0.08-0.12 s post-stimulus onset, corresponding to the M100 auditory evoked field, and plotted on a canonical MRI using FSL. Maps were thresholded at 75% of their maximum value, for illustrative purposes.

Next, a spatial filter was derived for an auditory cortex ROI and multiplied by the sensor-level data. The resulting virtual channel A1 data were averaged and plotted for each run (Fig. 6). The auditory ERF waveforms from this ROI were remarkably consistent across all three runs, with latency peaks corresponding to the M100 and M200 (Hari, 1990; Taulu & Hari, 2009). In participant 1 there was also an earlier peak, potentially corresponding to the M50 (Hari, 1990; Taulu & Hari, 2009).

**Fig. 6.**
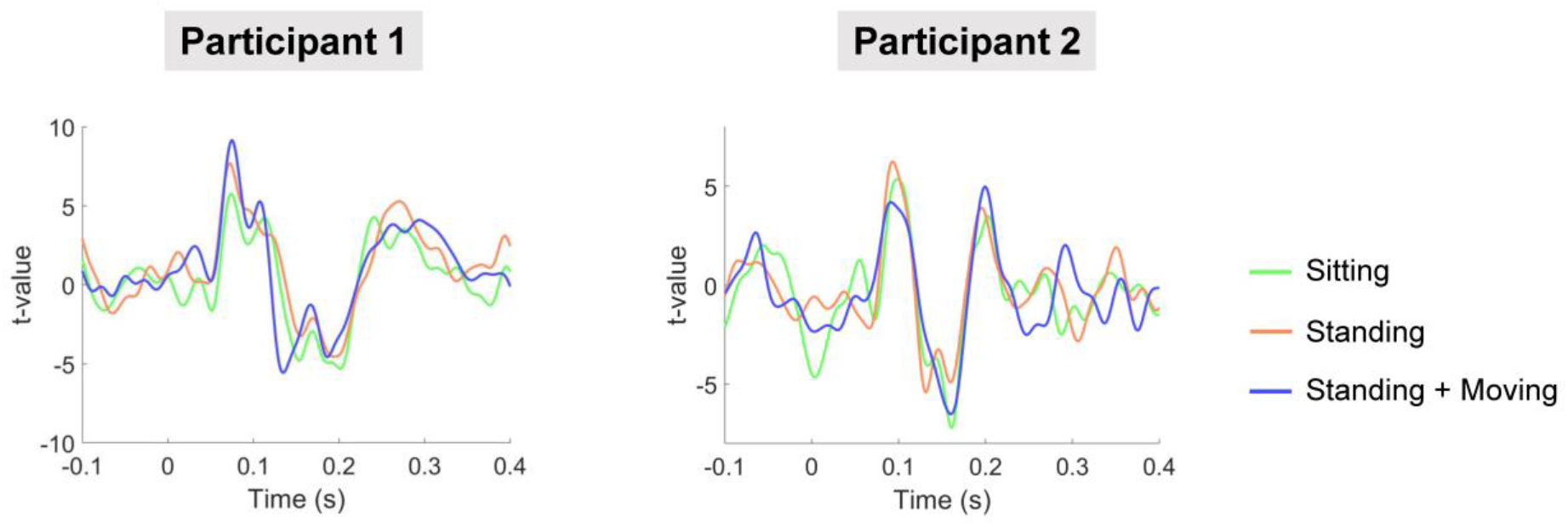
Evoked waveforms plotted from the auditory cortex ROI. Note the similarity in t-values across the sitting, standing, standing and moving runs for both participants.

### 3.4 Interference suppression

The use of interference suppression techniques were crucial to obtain the results presented above, especially when participants were standing up and moving their head. The raw data from run 3 contained mean trial-by-trial field changes of over 650 pT (see Supplementary Fig. S1), whereas AEFs are generally on the order of 50-150 fT. These large field changes are caused by the OPM sensors moving through field gradients in the MSR, which are particularly steep away from the centre of the room. However, it is worth noting that the relationship between the overall extent of movement and field change is often nonlinear (see Supplementary Fig. S5).

To characterise this interference and its removal, we calculated the power spectral density (PSD) between 2-10 Hz (below 2 Hz the data were cleaned using a high-pass filter). For the raw sensor-level data, the standing and standing and moving runs had higher PSD values below 6 Hz, compared to the sitting run (Fig. 7A). This demonstrates how low frequency artefacts are increased when participants are standing up, and especially when moving their head. The artefacts are particularly troublesome in this case because they overlap in the frequency domain with the neural signal of interest, namely the AEF (Hari, 1990). After pre-processing, the low-frequency artefacts are reduced but not eliminated for the standing and moving run (Fig. 7B). However, after beamforming, all three runs had equivalent PSD values from 2-10 Hz (Fig. 7C). Overall, we can see how movement-related artefacts below ∼6 Hz were progressively removed, first through the sensor-level pre-processing pipeline, and then through beamforming.

**Fig 7.**
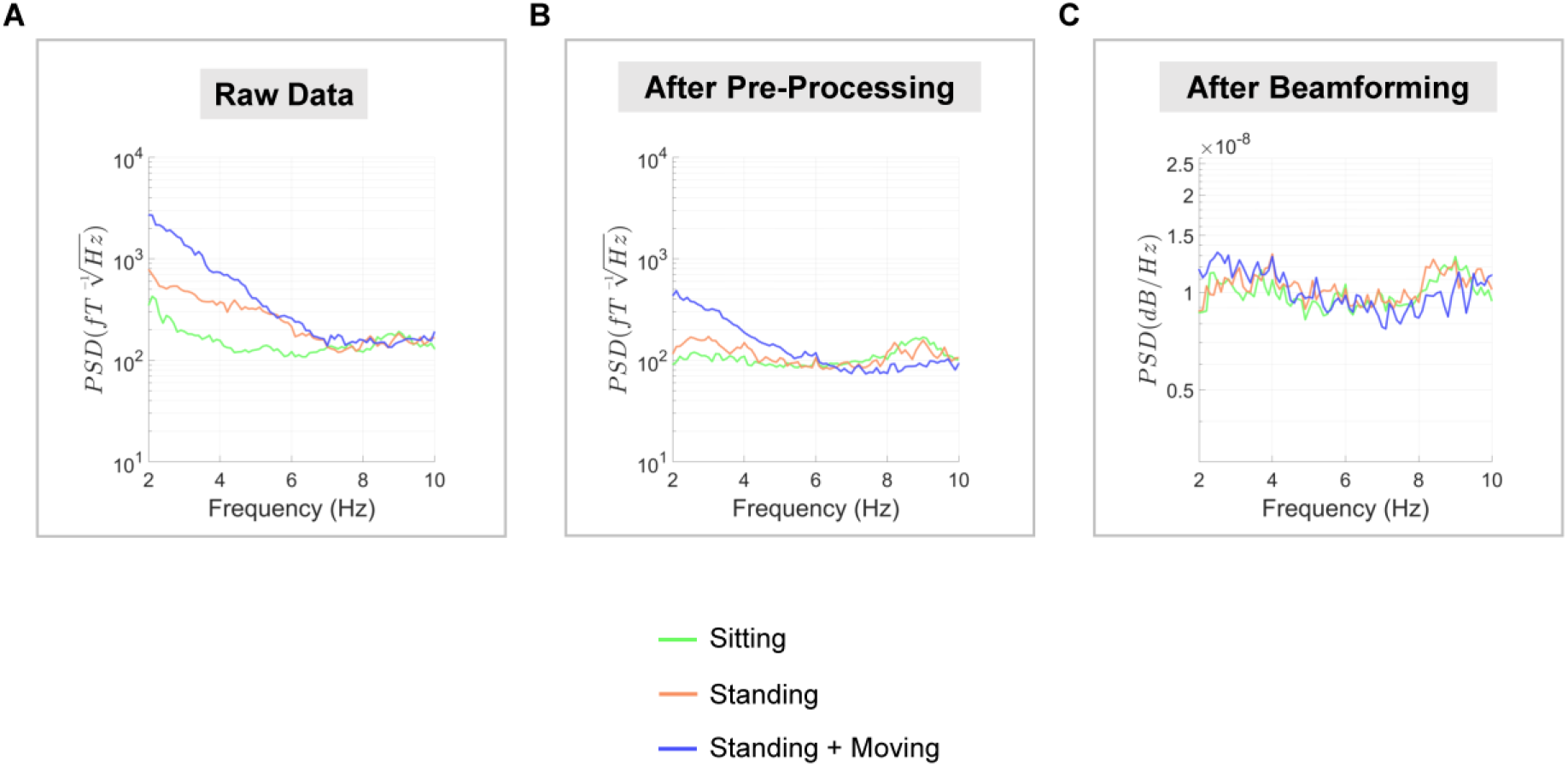
Removal of movement-related artefacts. **(A-B)** Power spectral density (PSD) was calculated for the OPM sensor with the greatest M100 response using both raw data and after pre-processing. **(C)** PSD was also calculated using the source-level auditory cortex ROI data. For all plots, PSD spectra are plotted between 2-10 Hz, and were averaged over the two participants.

To demonstrate the effectiveness of the LCMV beamformer for low-frequency interference suppression, we re-ran the analysis using varying levels of regularisation. This has the effect of making the beamformer less spatially-specific and therefore the spatial filtering properties will be reduced as regularisation increases. As expected, slowly increasing the regularisation from 0% to 1000% produced progressively less focal whole-brain localisation results (see Supplementary Fig. S6). In addition, the PSD values from the auditory cortex ROI showed that making the beamformer less spatially specific led to decreased suppression of low frequency movement artefacts (Supplementary Fig. S7).

Finally, to examine the impact of the movement data regression and homogenous-field correction (Tierney et al., 2021b) steps in greater detail, we re-ran our analysis pipeline for run 3 (standing and moving) using: (i) no movement data regression or homogenous-field correction; (ii) only movement data regression; (iii) only homogenous-field correction; and (iv) both movement data regression and homogenous-field correction. All other steps were the same across the analyses. PSD was calculated at the sensor-level using the channel with the largest M100 response (see Fig. 8, left). Results showed that the movement data regression step mainly reduced interference under ∼0.5Hz and had minimal impact at higher frequencies. In contrast, homogenous-field correction reduced interference across the spectrum from 0-10Hz. At the source level (Fig. 8, right), results showed that following beamforming, when data were processed with movement data regression and/or homogenous-field correction, they had very similar PSD values. Interestingly, processing without movement data regression or homogenous-field correction gave rise to slightly higher interference below 6Hz. In terms of the impact on AEFs, homogenous-field correction had the most impact at the sensor-level, increasing t-values when used alone and in combination with movement data regression (see Supplementary Fig. S8). Following beamforming, all data showed remarkably similar evoked waveforms, irrespective of whether movement data regression and/or homogenous-field correction was applied (see Supplementary Fig. S9), further highlighting the interference suppression quality of the beamformer.

**Fig 8.**
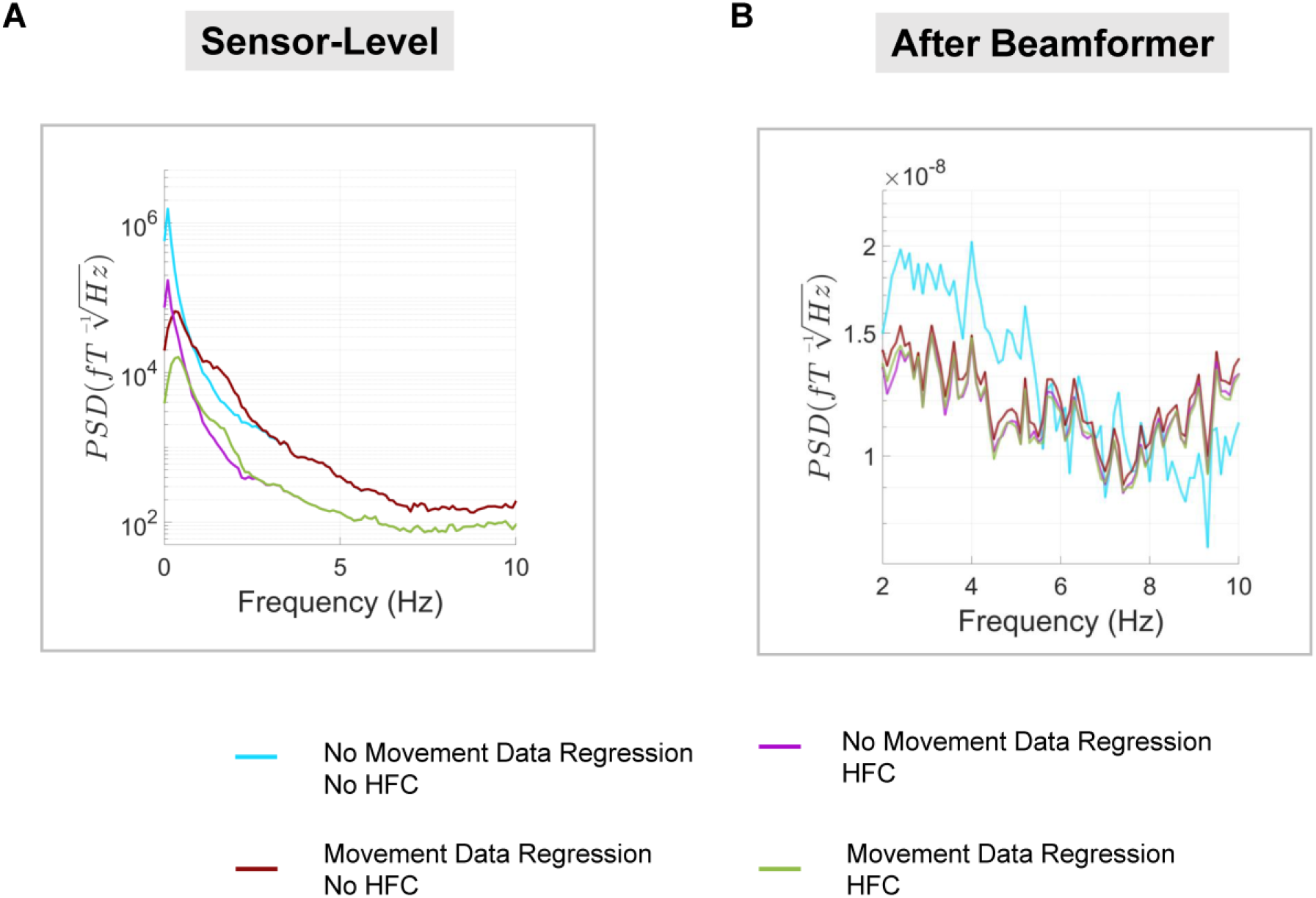
Investigating sensor-level pre-processing steps. **(A)** Power spectral density (PSD) was calculated for the OPM sensor with the greatest M100 response. PSD was plotted separately for sensor-level data with/without the movement data regression step, and with/without homogenous-field correction. **(B)** PSD was calculated in the same way using the source-level auditory cortex ROI data. For all plots, PSD spectra were averaged over the two participants.

## 4. Discussion

We set out to demonstrate, as a proof-of-principle, that neuromagnetic fields can be successfully recorded with OP-MEG while participants are standing up and making natural and continuous head movements. To do this, we measured AEFs across three scanning runs while two participants were: (i) sitting down with their heads stationary, (ii) standing up with their heads stationary, and (iii) standing up and deliberately moving their heads continuously, with large translations and rotations, tracked using a six-camera motion capture system. Across all three runs we found reliable AEFs (M100) at the source level localised to primary auditory cortex.

Previous OP-MEG studies have successfully measured neuromagnetic fields during ∼20 cm translations and ∼30° rotations, across the course of an experiment while participants were sitting near the centre of an MSR (Boto et al., 2018; Hill et al., 2019; Holmes et al., 2018; Roberts et al., 2019). Extending these findings, we have shown here that OP-MEG can be used to successfully detect AEFs while two participants were standing up and making natural movements of their heads of a much greater magnitude than those reported previously – maximum translation >100 cm; maximum rotation >190° – over the course of an experiment. It is also important to emphasise that head movements in run 3 of our study were continuous, with the two participants moving their heads on average ∼5 cm during each 0.5s-long trial.

The successful measurement of neural activity in standing, moving participants is an important technical milestone, given that residual field gradients inside MSRs are generally steeper away from the centre of the room, exacerbating low frequency movement related artefacts. In our study, the raw data contained field changes on the order of ∼650 pT per trial when the two participants were standing and moving (see Supplementary Fig. S3, S5). Several steps were taken to alleviate these artefacts. The remnant background field was reduced using a custom-designed MSR with degaussing coils. Second generation OPM sensors also have on-board coils designed to cancel static fields inside the OPM cell (Osborne et al., 2018). In addition, a series of offline interference suppression techniques were used. The motion capture data (translations and rotations) were included in a linear regression to remove OP-MEG data covarying with participant movement. This was successful in removing interference under 0.5Hz, when the two participants were standing and moving (run 3). However, when the two participants were sitting or standing still, the movement data regression step actually introduced noise into the data. Future work should focus on reducing noise from the motion-tracking recordings. The next pre-processing step involved modelling external interference as a homogenous field and removing this from the data (Tierney et al., 2021b). This step helped to reduce interference across the frequency spectrum, and increased the detectability of AEFs at the sensor-level. Future studies could explore the option of modelling the interfering field gradients from participant movement and incorporating additional terms into the basis set based on higher-order spherical harmonic expansions of the data. Alternatively, field-mapping approaches could be used to remove movement-related interference (Mellor et al., 2021). Next, a 2 Hz high-pass filter was used to attenuate low-frequency drifts in the data that are known to overlap substantially with movement-related artefacts. As a final interference suppression step, we used an LCMV beamformer for source localisation, which reduces the influence of non-neural signals in the data via spatial filtering (Boto et al., 2016; Hillebrand & Barnes, 2005). By combining all of these steps, movement-related artefacts were reduced (see Fig. 7), and consistent, interpretable AEF waveforms were measured, irrespective of whether the participant was sitting still, standing still, or standing and moving their head (Fig. 5, 6).

If the remnant background field within the MSR can be kept as low as possible, and OPM sensors can be maintained within their dynamic range, even larger translations or rotations of the head may be possible in future experiments. It should be noted, however, that the range of movements possible with current OP-MEG technology is not infinite. Where translations and rotations of the sensors against a static field produce changes over ∼5 nT (especially common during large and rapid rotations of the head), the dynamic range of the current second generation QuSpin sensors will be exceeded. Where more substantial rotations and/or faster head movements need to be incorporated into experimental designs, the remnant background field in the MSR could be further reduced using external nulling coils (e.g. Holmes et al., 2018, 2019). Alternatively, OPMs could be adapted to operate in a ‘closed-loop’ mode, where the remnant magnetic fields are continuously modelled (for example, using a spherical harmonic field model (as in Mellor et al., 2021) and cancelled (Fourcault et al., 2021; Nardelli et al., 2020).

In spite of the pre-processing steps, the sensor-level data from the standing and moving run were contaminated by low-frequency movement artefacts (Fig. 7B, Supplementary Fig. S4). However, upon utilisation of an LCMV beamformer, the data across the three runs were remarkably consistent, both in terms of M100 localisation and the ERF waveforms in primary auditory cortex. The beamformer was even able to successfully remove most of the interference when the sensor-level movement data regression and homogenous-field correction steps were omitted (Supplementary Fig. S9). The use of beamforming for interference suppression is well established (Fatima et al., 2013; Hillebrand & Barnes, 2005). In much the same way as conventional frequency filters select signals only within a specified temporal range, the beamformer acts as a spatial filter to select only signals from specified spatial locations (Adjamian et al., 2009). Signals arising from outside the brain (e.g. movement artefacts, cardiac activity and other environmental noise) are removed from the data. In support of this, we showed (Supplementary Fig. S6, S7) that by making the beamformer less spatially-specific (by increasing the beamformer regularisation parameter), the suppression of low-frequency movement artefacts was compromised (Litvak et al., 2010). Brookes et al. (2021) also recently showed how triaxial OPM sensor arrays can theoretically improve the interference suppression performance of beamformers even further.

The reliance of our results on the spatial filtering properties of beamformers has several implications for future OP-MEG experiments. Where OP-MEG data are required to be interpreted by clinicians at the sensor-level (e.g. for epilepsy pre-surgical evaluation), experimenters could project the cleaned source-level data back to the sensor-space using lead-field information. In situations where the exact neural signals of interest are unknown, an atlas-based, broadband beamforming approach could be used in combination with robust statistics to control for multiple comparisons over space and time (Maris & Oostenveld, 2007; Nichols & Holmes, 2002).

Considering next the auditory system, our results show that OP-MEG can be used to measure AEFs using 70 ms-long tones, replicating several previous stationary, seated OP-MEG studies (Borna et al., 2017; 2020; Kowalczyk et al., 2021). Given their reliability (Hari, 1990), AEFs will continue to be a crucial benchmarking tool for OP-MEG development. Here we showed that the localisation, latency and amplitude of AEFs can be measured *while* participants are standing and moving naturally within an MSR. The successful measurement of evoked magnetic fields during standing accompanied by substantial participant movement is a notable result, given the considerable overlap in the frequency domain between these evoked responses (2-40 Hz) and movement artefacts (<6 Hz).

More generally, our results show that OP-MEG neuromagnetic measurements are now feasible in participants who are less able to keep still, such as young children (Hill et al., 2019). This will facilitate advances in developmental neuroscience and will increase our understanding of conditions like Autism Spectrum Disorder (Kessler et al., 2016), which is associated with a variety of differences in early auditory and language processing (Roberts et al., 2011; Rojas & Wilson, 2014; Seymour et al., 2020). Clinically, OP-MEG is well-placed to assist in the detection and localisation of epileptic spikes for pre-surgical evaluation (Vivekananda et al., 2020), and the mapping of eloquent cortex in younger children (Tierney et al., 2018). From a neuroscientific perspective, our results open up exciting avenues for experimenters wishing to incorporate large natural movements from a standing position into neuroimaging paradigms, while also measuring magnetic fields from the brain with high temporal and spatial resolution. This could include natural interactions with objects or touch-screens (Jungnickel & Gramann, 2016), and two-person (e.g. parent-child) social neuroscience tasks. OP-MEG has also been used with rudimentary virtual reality technology (Roberts et al., 2019). Combining OP-MEG with more complex and realistic virtual reality environments, as well as MEG-compatible treadmills, could provide novel opportunities for cognitive neuroscientists interested in understanding the neural basis of, for example, the experiencing of real-life events and how such episodes are encoded and processed by the human brain.

In summary, here we collected neural data using a 45 sensor (90 channel) OPM array. Through a combination of passive magnetic shielding and post-hoc data processing, auditory evoked fields were successfully measured while two participants were standing up and making large, continuous movements of the head.

## Supplementary materials

Supplementary materials are available for this article.

## Declaration of interest

This work was partly funded by a Wellcome Collaborative Award that involves a collaboration agreement with QuSpin Inc.

## Data and code availability

The data that support the findings of this study are available from Zenodo: (https://doi.org/10.5281/zenodo.5172689), under an Attribution-NonCommercial 3.0 Unported license. Analysis code is openly available on GitHub:

(https://github.com/FIL-OPMEG/movement_auditory_ERF).

## Funding information

This research was supported by a Wellcome Principal Research Fellowship to E.A.M. (210567/Z/18/Z), a Wellcome Collaborative Award (203257/Z/16/Z), a Wellcome Centre Award (203147/Z/16/Z), the EPSRC-funded UCL Centre for Doctoral Training in Medical Imaging (EP/L016478/1), the Department of Health’s NIHR-funded Biomedical Research Centre at University College London Hospitals, and EPSRC (EP/T001046/1) funding from the Quantum Technology hub in sensing and timing (sub-award QTPRF02).

This research was funded in whole, or in part, by Wellcome (Grant numbers: 210567/Z/18/Z; 203257/Z/16/Z; 203147/Z/16/Z). For the purpose of Open Access, the authors have applied a CC BY public copyright licence to any Author Accepted Manuscript version arising from this submission.

## CRediT authorship contribution statement

**Robert A. Seymour:** Conceptualisation, Methodology, Software, Investigation, Formal Analysis, Writing – Original Draft; **Nicholas Alexander:** Conceptualisation, Methodology, Software, Investigation, Formal Analysis, Writing – Original Draft; **Stephanie Mellor:** Software, Resources, Writing – Review and Editing; **George C. O’Neill:** Software, Resources, Writing – Review and Editing; **Tim M. Tierney:** Software, Resources, Writing – Review and Editing; **Gareth R. Barnes:** Software, Writing – Review and Editing; **Eleanor A. Maguire:** Conceptualisation, Methodology, Supervision, Funding Acquisition; Writing – Review and Editing.

## Acknowledgements

Thanks to Vladimir Litvak and Sven Bestmann for valuable discussions, David Bradbury for imaging support, Vishal Shah at QuSpin Inc. for technical assistance, and Mark Lim at Chalk Studios for help with scanner-cast design and construction. Thanks also to Matthew Brookes, Richard Bowtell and their team at the University of Nottingham for useful interactions.

## Supplementary Materials

**Supplementary Table S1.**
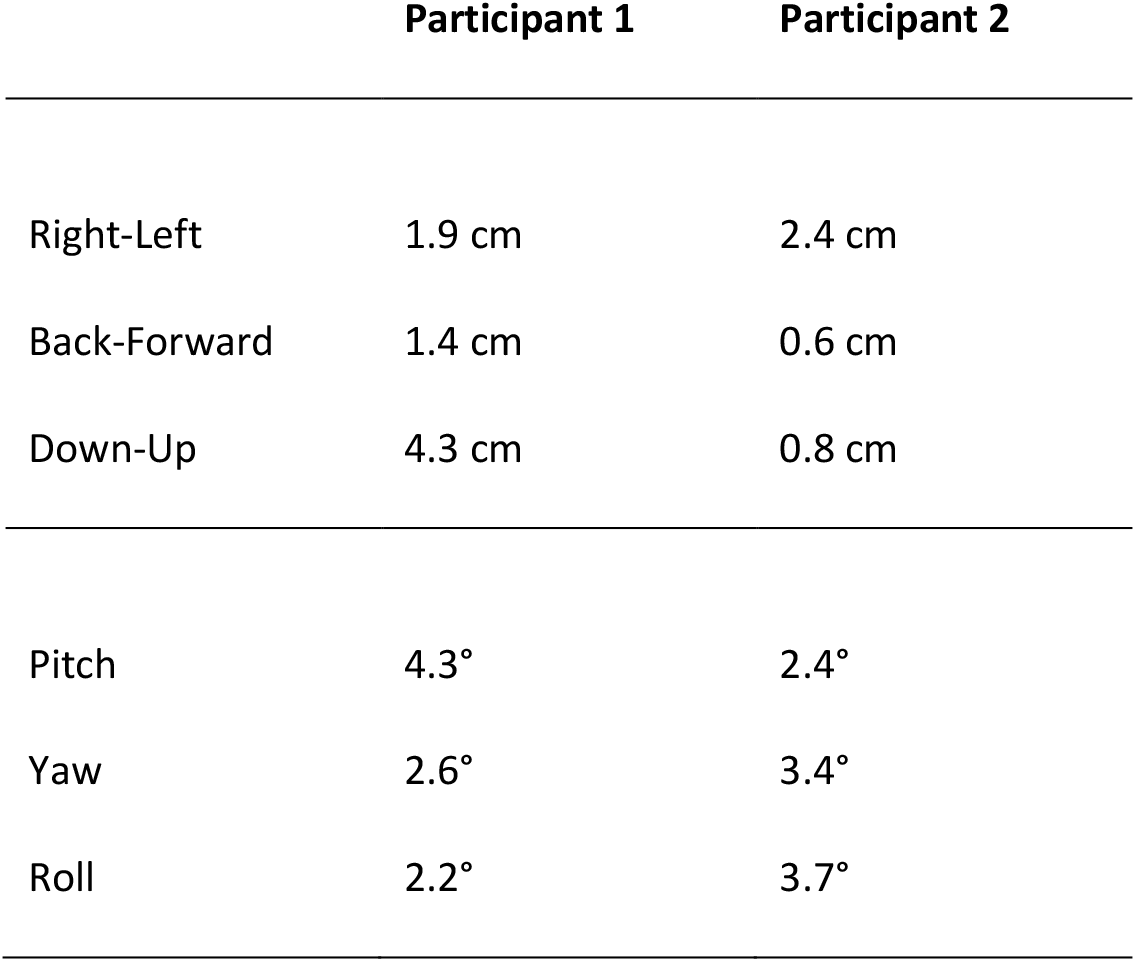
Rigid body data. Range of values across the sitting still condition (run 1), for each of the six degrees of freedom.

**Supplementary Table S2.** Rigid body data. Range of values across the standing still condition (run 2), for each of the six degrees of freedom.

|  | Participant 1 | Participant 2 |
| --- | --- | --- |
| Right-Left | 2.5 cm | 6.5 cm |
| Back-Forward | 4.4 cm | 2.8 cm |
| Down-Up | 6.7 cm | 2.7 cm |
| Pitch | 3.7° | 12.5° |
| Yaw | 4.4° | 5.9° |
| Roll | 3.5° | 0.4° |

**Supplementary Fig. S1.**
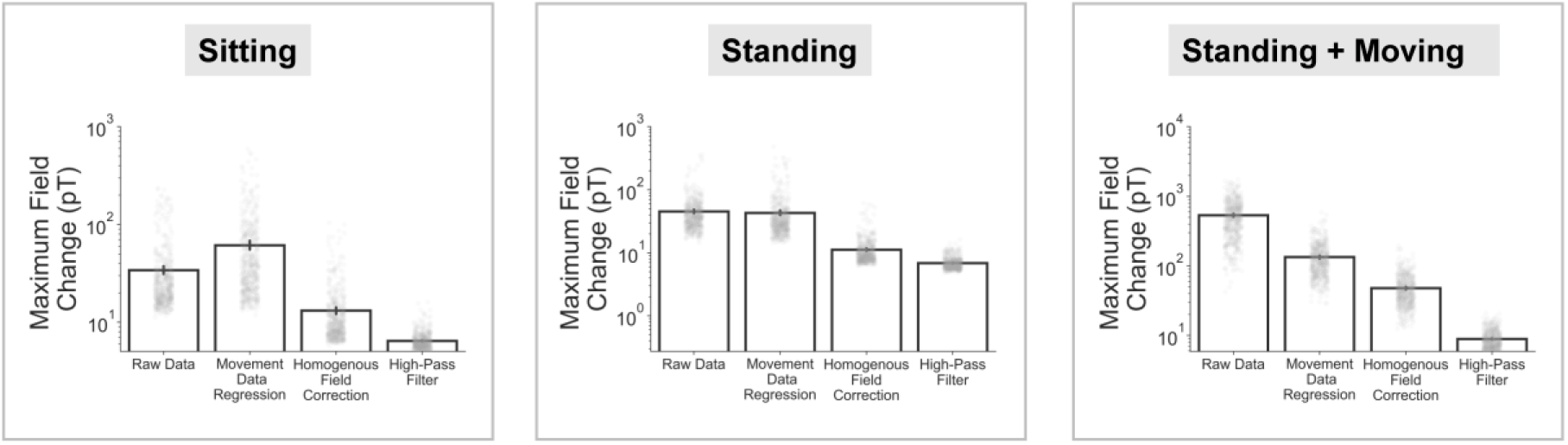
For each run the maximum change in magnetic field was calculated for each trial, using the raw data and after each of the pre-processing steps (data are averaged across the two participants). Individual data points (corresponding to each trial) are plotted in grey.

**Supplementary Fig. S2.**
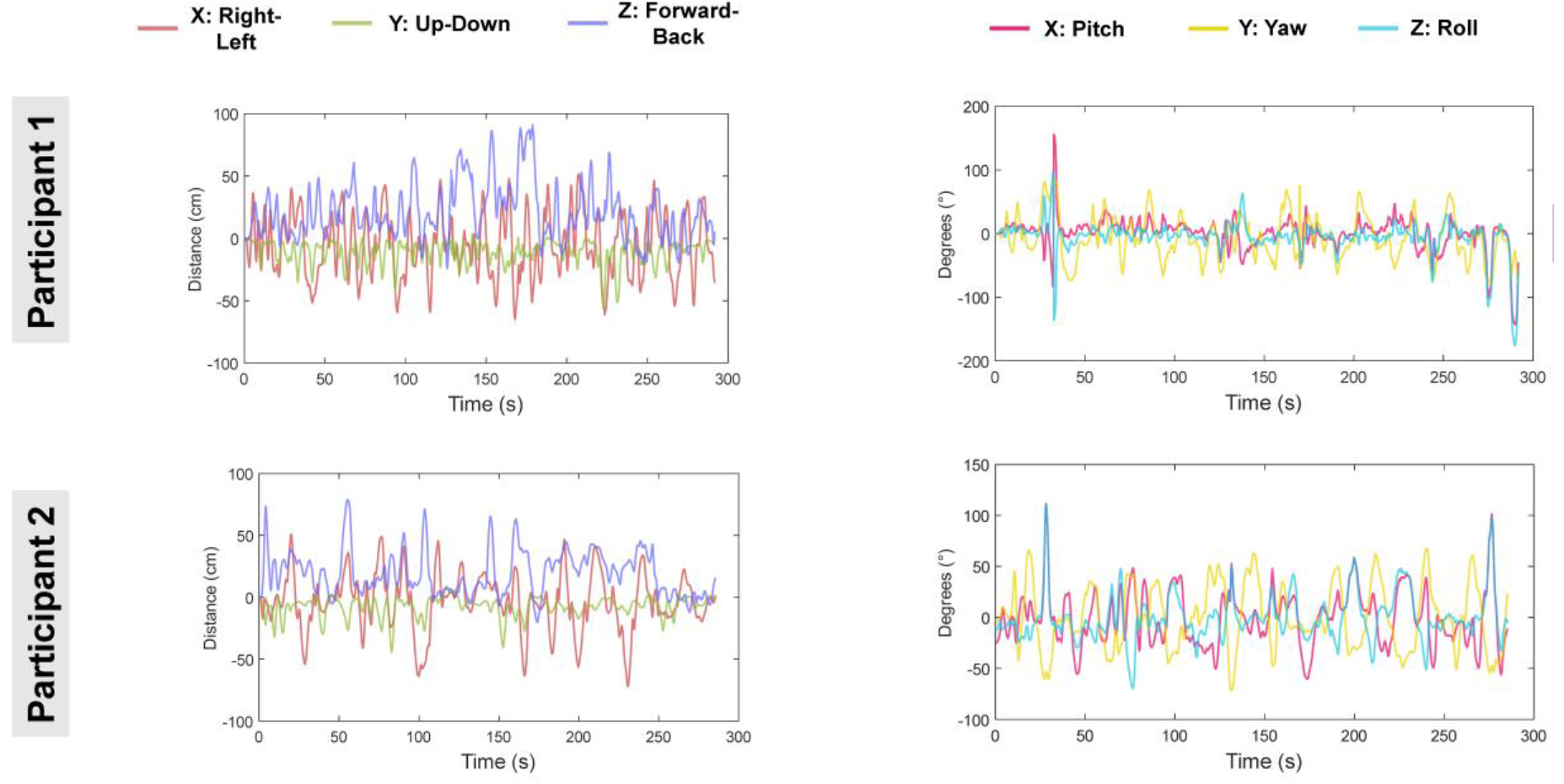
For run 3 (standing and moving), continuous rigid body data were plotted over time for both participants. Left panels = translations, right panel = rotations. Note the continuous nature of the movements over the course of the entire auditory experiment.

**Supplementary Fig. S3.**
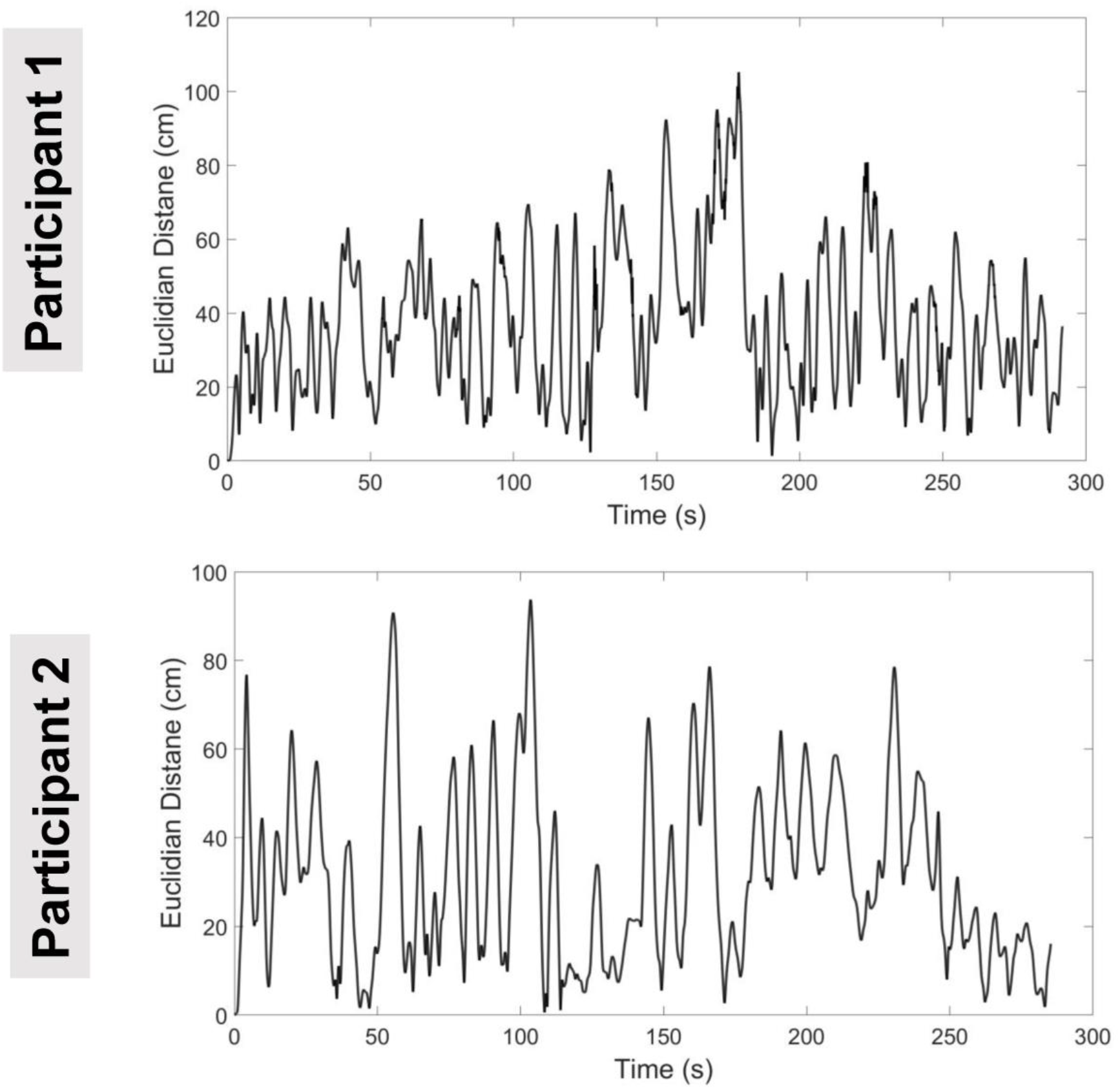
Continuous rigid body Euclidian distances from the start point of run 3, were plotted for both participants. Note the continuous nature of the movements over the course of the entire auditory experiment.

**Supplementary Fig. S4.**
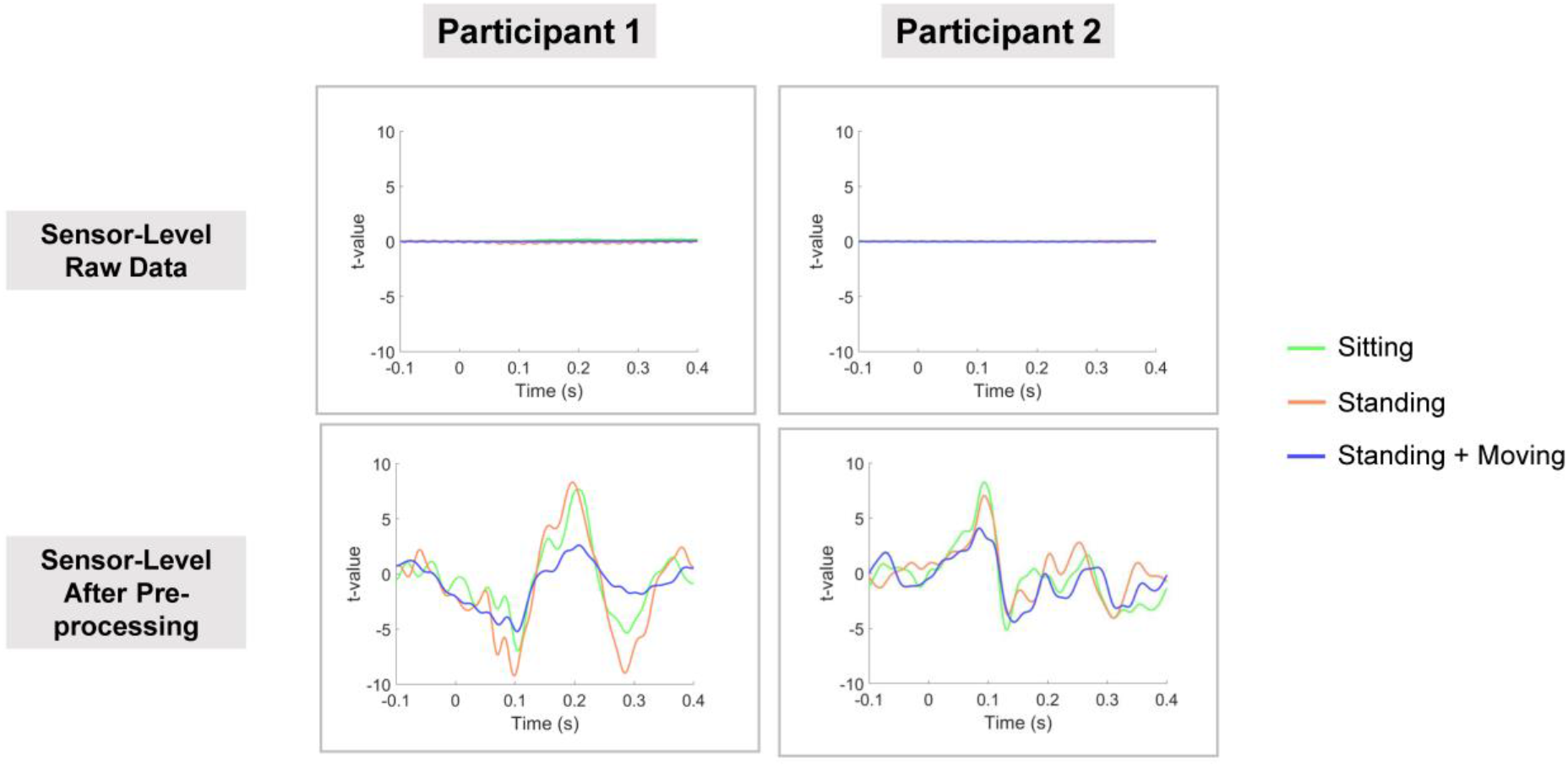
For each participant, auditory ERF t-values were calculated for the OPM sensor with the greatest M100 response using the raw data and after pre-processing. The OPM sensor was the same across all three runs (Participant 1: N3-TAN; Participant 2: 1A-TAN) and was located approximately over the left superior temporal lobe. The pipeline for pre-processing was almost identical across three runs, except that the movement data regression step was applied to only the standing and moving (run 3) data. The raw data showed no clear ERF for any run. After pre-processing (bottom panel), auditory ERFs were very similar for the sitting (green line) and standing (orange) runs, but reduced for the standing and moving run (blue line).

**Supplementary Fig. S5.**
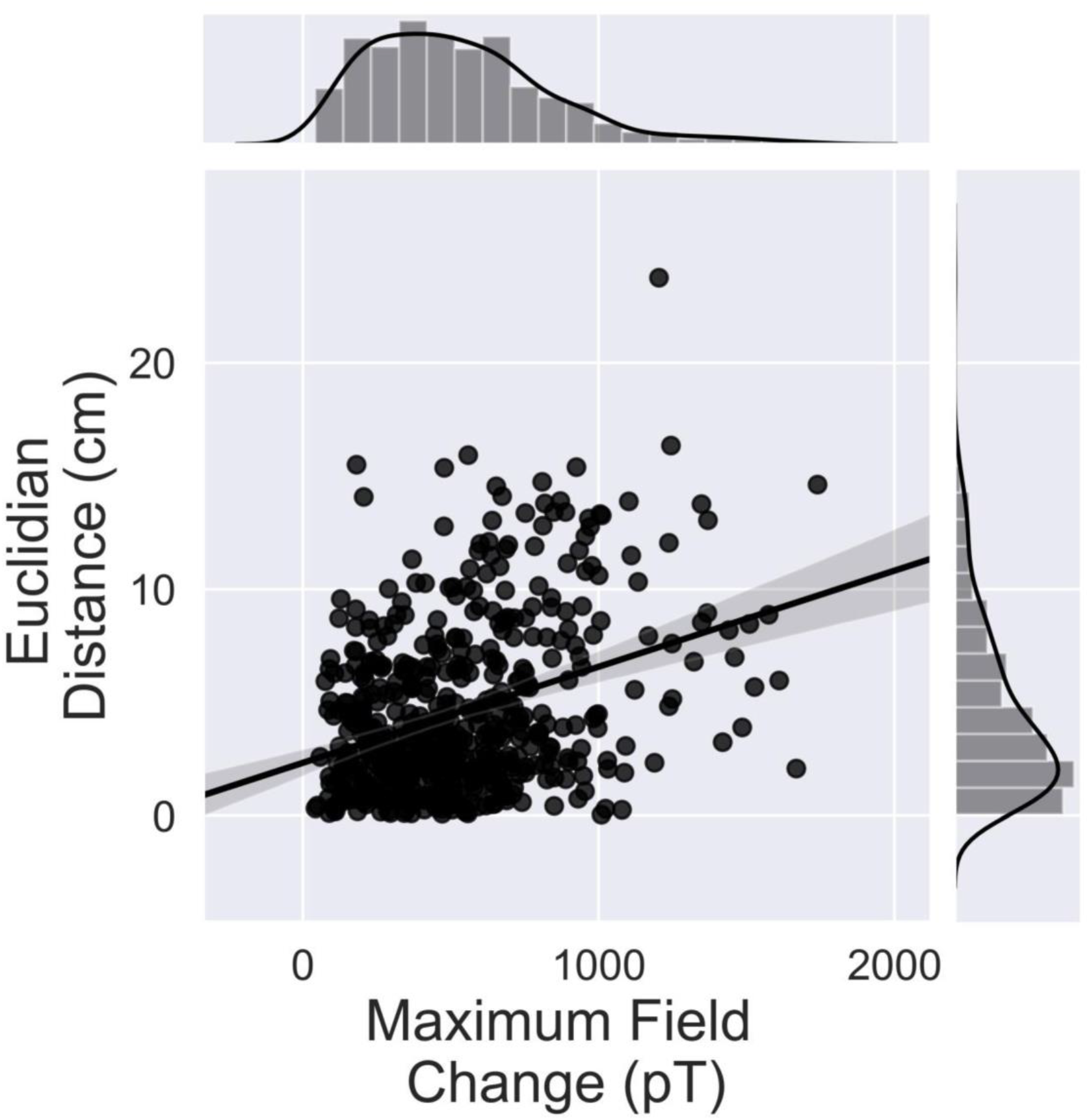
For Participant 2, a scatter-plot with regression line was produced to show the relationship between Euclidian distance moved per trial and the maximum field change. Using a Pearson’s correlation, we found no statistically significant relationship between the two, r=0.052, p=0.237.

**Supplementary Fig. S6.**
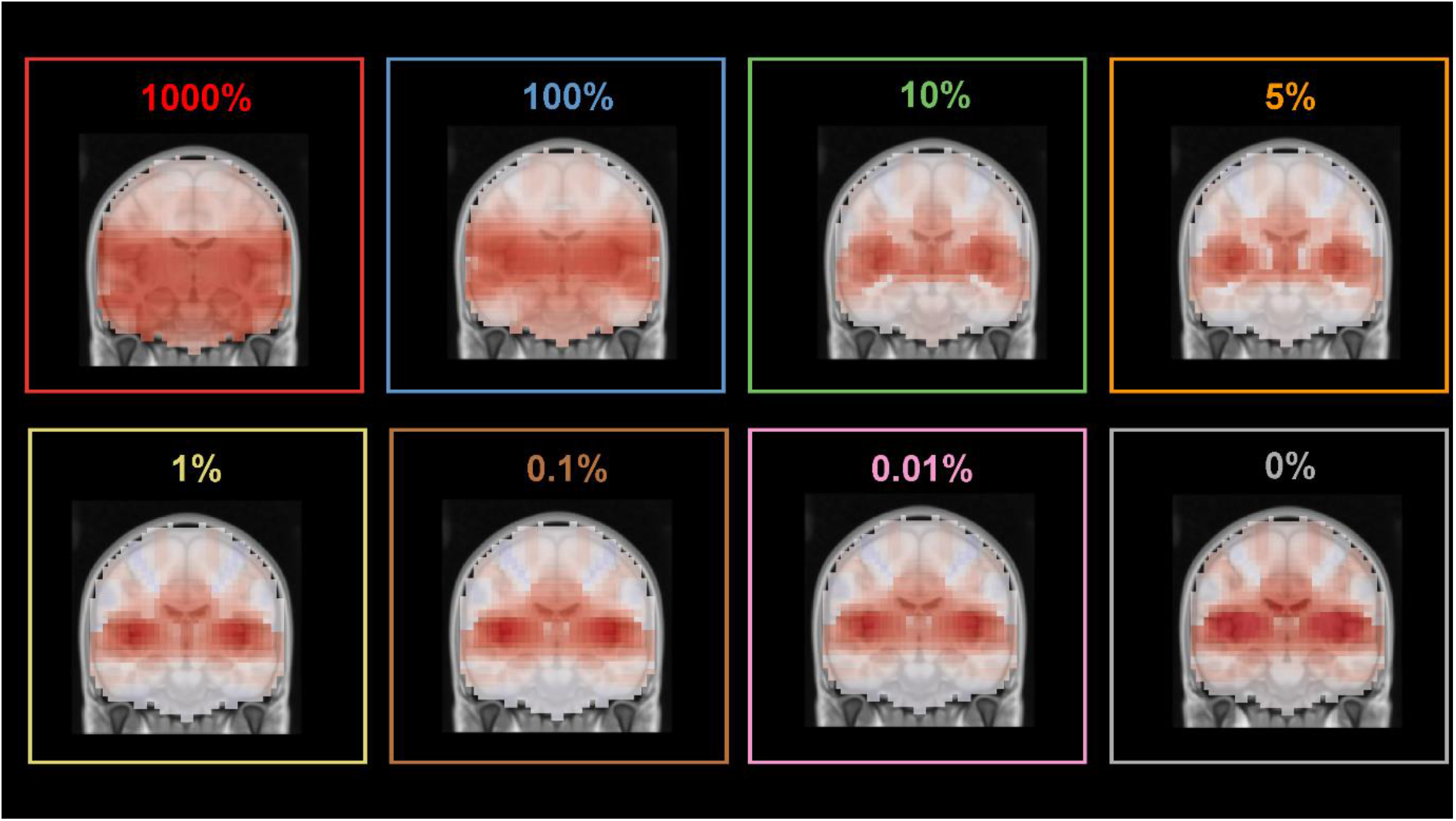
For participant 2, run 3 (standing and moving), whole-brain source localisation was repeated. The LCMV beamformer regularisation parameter was gradually increased from 0% to 1000%. Note the less focal whole-brain Neural Activity Index maps at 1000% and 100%.

**Supplementary Fig. S7.**
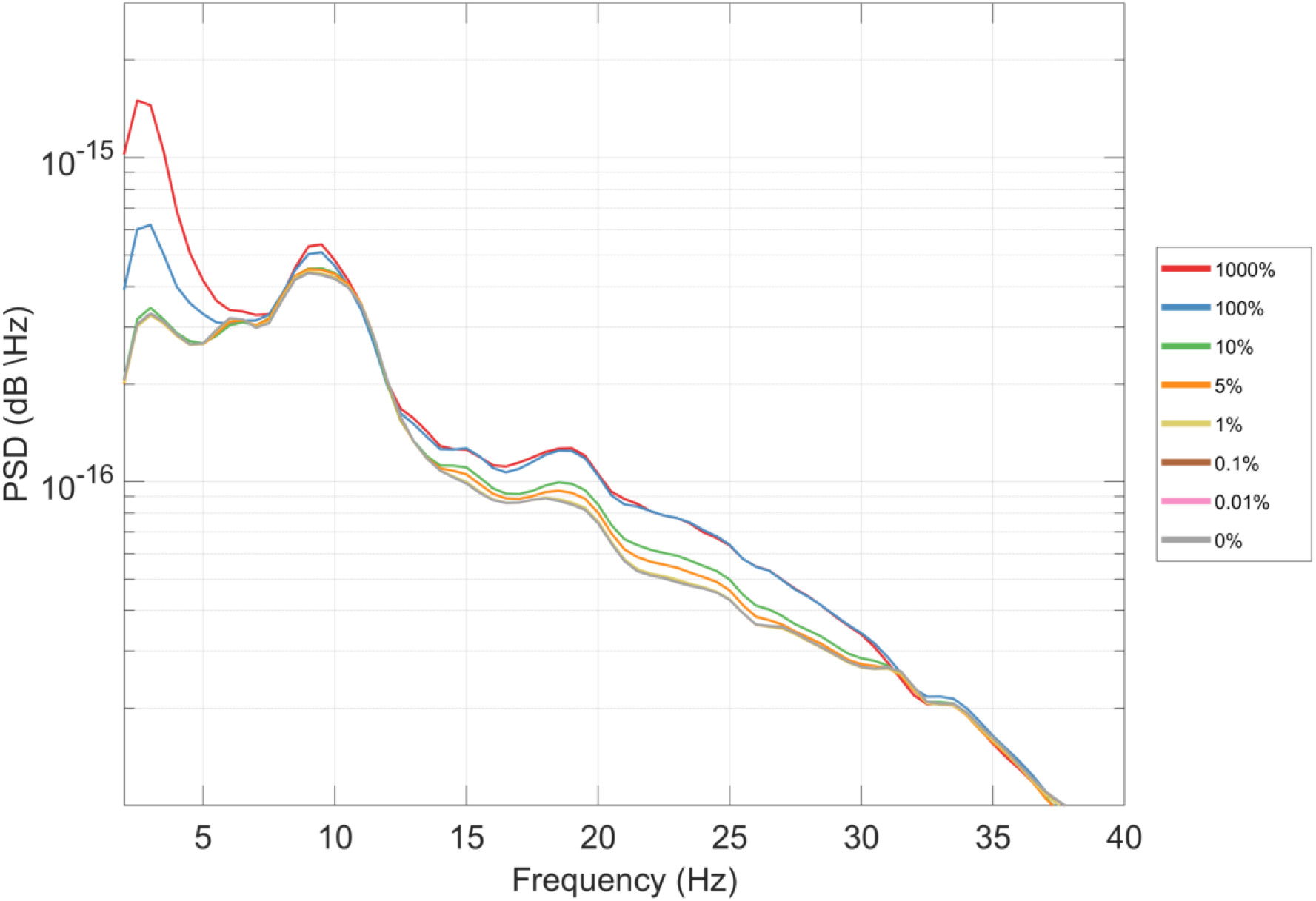
For participant 2, power spectral density (PSD) was calculated using Welch’s method using data from the auditory cortex region of interest. The LCMV beamformer regularisation parameter was gradually varied between 0% and 1000%. Note the gradual reduction in low-frequency interference suppression as regularisation increased (especially between 2-6 Hz). Also note that the lines for 0%, 0.01% and 0.1% are virtually indistinguishable on the graph, showing how similar the PSD values were between these three levels of regularisation.

**Supplementary Fig. S8.**
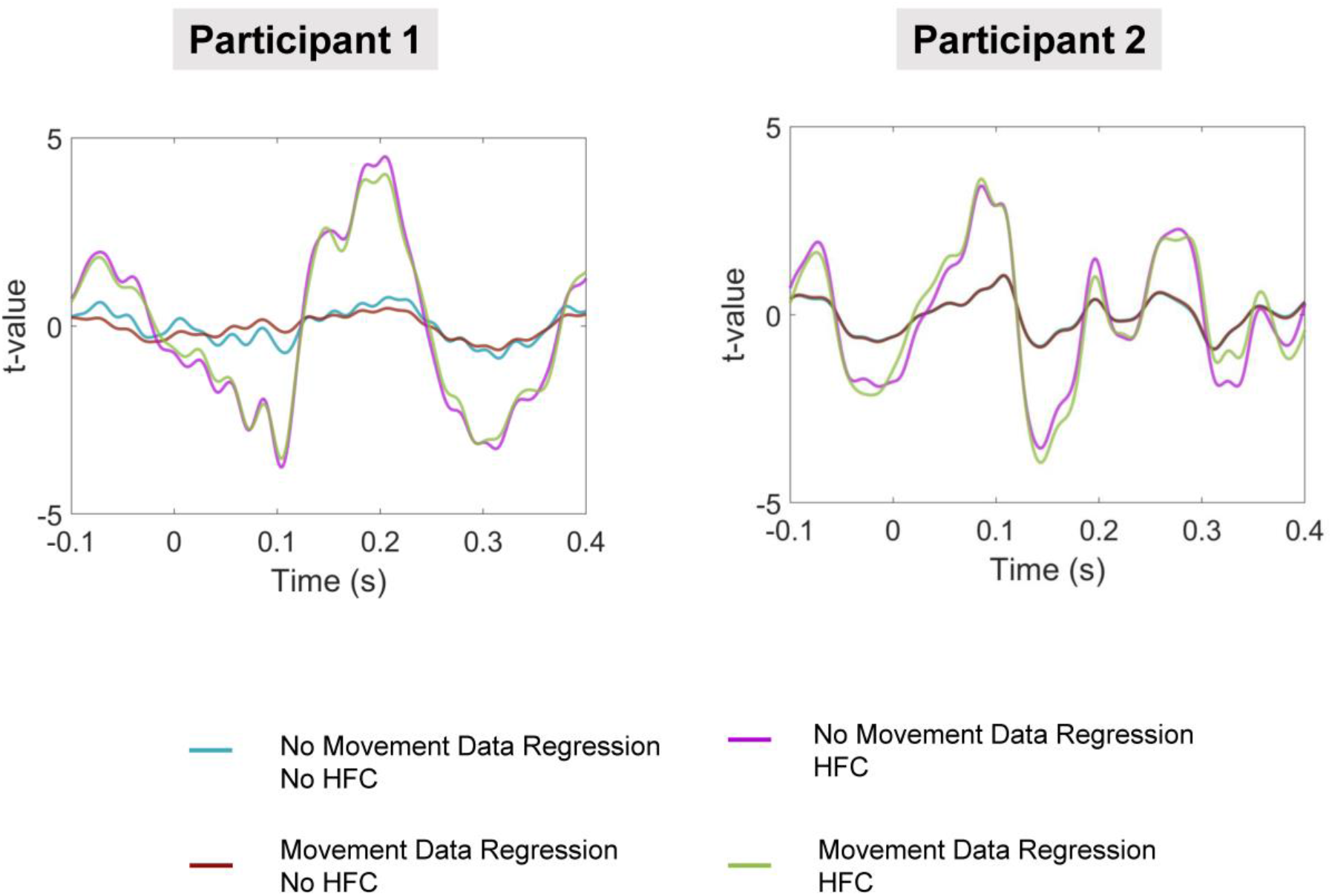
For each participant, auditory ERF t-values were calculated for the OPM sensor with the greatest M100 response using data from run 3 (standing and moving), located approximately over the left superior temporal lobe. The different coloured lines correspond to various permutations of pre-processing: with or without movement data regression, and with or without homogenous-field correction.

**Supplementary Fig. S9.**
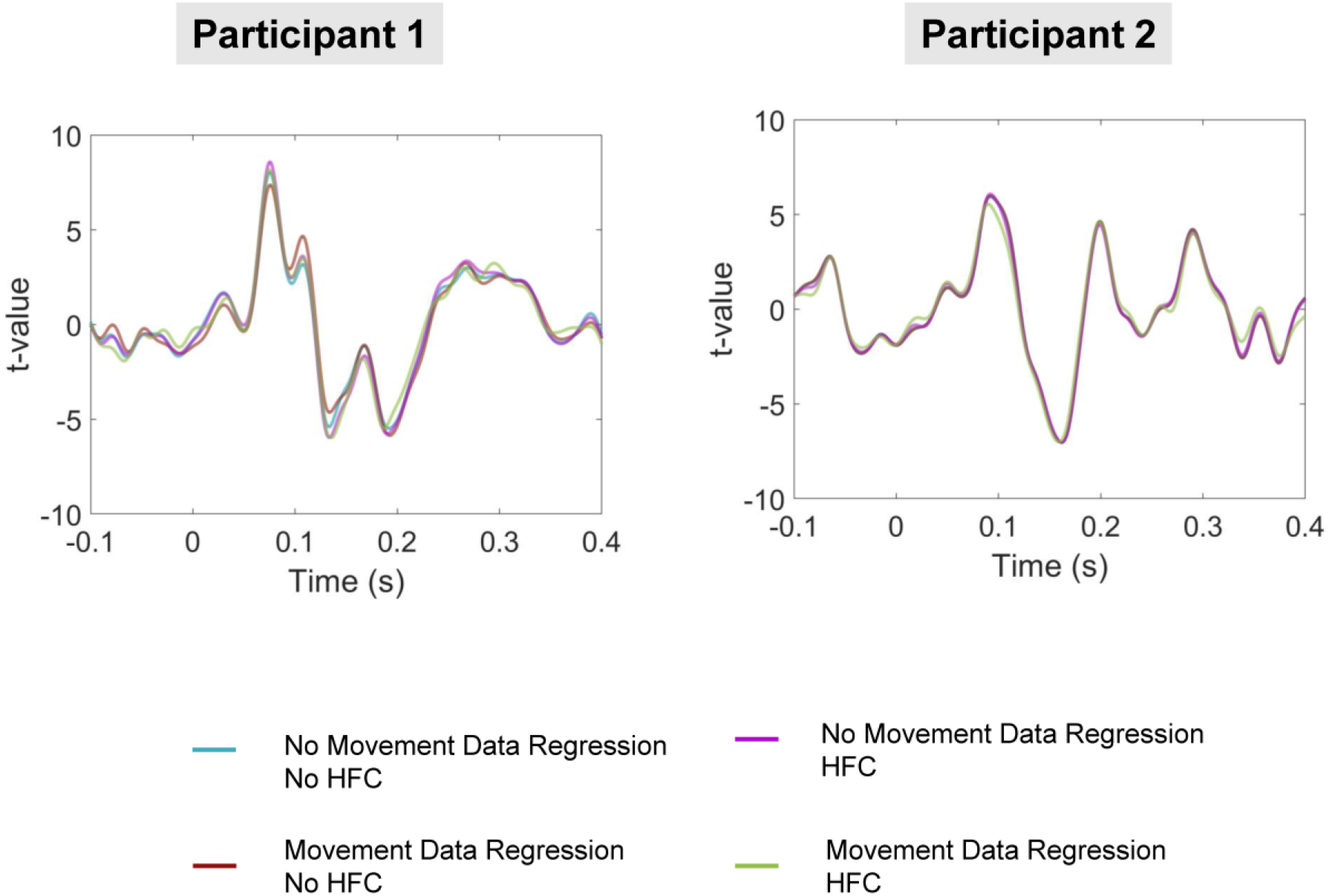
For each participant, t-values were calculated for the auditory cortex region of interest, using data from run 3 (standing and moving). The different coloured lines correspond to various permutations of pre-processing: with or without movement data regression, and with or without homogenous-field correction. Note that the lines are largely overlapping.

## References

Adjamian, P., Worthen, S. F., Hillebrand, A., Furlong, P. L., Chizh, B. A., Hobson, A. R., Aziz, Q., & Barnes, G. R. (2009). Effective electromagnetic noise cancellation with beamformers and synthetic gradiometry in shielded and partly shielded environments. Journal of Neuroscience Methods, 178(1), 120–127.

Altarev, I., Fierlinger, P., Lins, T., Marino, M. G., Nießen, B., Petzoldt, G., Reisner, M., Stuiber, S., Sturm, M., Taggart Singh, J., Taubenheim, B., Rohrer, H. K., & Schläpfer, U. (2015). Minimizing magnetic fields for precision experiments. Journal of Applied Physics, 117(23), 233903.

Baillet, S. (2017). Magnetoencephalography for brain electrophysiology and imaging. Nature Neuroscience, 20(3), 327–339.

Barry, D. N., Tierney, T. M., Holmes, N., Boto, E., Roberts, G., Leggett, J., Bowtell, R., Brookes, M. J., Barnes, G. R., & Maguire, E. A. (2019). Imaging the human hippocampus with optically-pumped magnetoencephalography. NeuroImage, 203, 116192.

Borna, A., Carter, T. R., Colombo, A. P., Jau, Y.-Y., McKay, J., Weisend, M., Taulu, S., Stephen, J. M., & Schwindt, P. D. D. (2020). Non-Invasive Functional-Brain-Imaging with an OPM-based Magnetoencephalography System. PLOS ONE, 15(1), e0227684.

Borna, A., Carter, T. R., Goldberg, J. D., Colombo, A. P., Jau, Y.-Y., Berry, C., McKay, J., Stephen, J., Weisend, M., & Schwindt, P. D. (2017). A 20-channel magnetoencephalography system based on optically pumped magnetometers. Physics in Medicine & Biology, 62(23), 8909.

Boto, E., Bowtell, R., Krüger, P., Fromhold, T. M., Morris, P. G., Meyer, S. S., Barnes, G. R., & Brookes, M. J. (2016). On the potential of a new generation of magnetometers for MEG: A beamformer simulation study. PloS One, 11(8), e0157655.

Boto, E., Hill, R. M., Rea, M., Holmes, N., Seedat, Z. A., Leggett, J., Shah, V., Osborne, J., Bowtell, R., & Brookes, M. J. (2021). Measuring functional connectivity with wearable MEG. NeuroImage, 117815.

Boto, E., Holmes, N., Leggett, J., Roberts, G., Shah, V., Meyer, S. S., Muñoz, L. D., Mullinger, K. J., Tierney, T. M., & Bestmann, S. (2018). Moving magnetoencephalography towards real-world applications with a wearable system. Nature, 555(7698), 657–661.

Boto, E., Meyer, S. S., Shah, V., Alem, O., Knappe, S., Kruger, P., Fromhold, T. M., Lim, M., Glover, P. M., & Morris, P. G. (2017). A new generation of magnetoencephalography: Room temperature measurements using optically-pumped magnetometers. NeuroImage, 149, 404–414.

Brookes, M. J., Boto, E., Rea, M., Shah, V., Osborne, J., Holmes, N., Hill, R. M., Leggett, J., Rhodes, N., & Bowtell, R. (2021). Theoretical advantages of a triaxial optically pumped magnetometer magnetoencephalography system. NeuroImage, 118025.

Brookes, M. J., Stevenson, C. M., Barnes, G. R., Hillebrand, A., Simpson, M. I. G., Francis, S. T., & Morris, P. G. (2007). Beamformer reconstruction of correlated sources using a modified source model. NeuroImage, 34(4), 1454–1465.

Brookes, M. J., Vrba, J., Robinson, S. E., Stevenson, C. M., Peters, A. M., Barnes, G. R., Hillebrand, A., & Morris, P. G. (2008). Optimising experimental design for MEG beamformer imaging. Neuroimage, 39(4), 1788–1802.

de Cheveigné, A., & Nelken, I. (2019). Filters: When, Why, and How (Not) to Use Them. Neuron, 102(2), 280–293.

Fatima, Z., Quraan, M. A., Kovacevic, N., & McIntosh, A. R. (2013). ICA-based artifact correction improves spatial localization of adaptive spatial filters in MEG. NeuroImage, 78, 284–294.

Fourcault, W., Romain, R., Le Gal, G., Bertrand, F., Josselin, V., Le Prado, M., Labyt, E., & Palacios-Laloy, (2021). Helium-4 magnetometers for room-temperature biomedical imaging: Toward collective operation and photon-noise limited sensitivity. Optics Express, 29(10), 14467–14475.

Garrido, M. I., Friston, K. J., Kiebel, S. J., Stephan, K. E., Baldeweg, T., & Kilner, J. M. (2008). The functional anatomy of the MMN: A DCM study of the roving paradigm. Neuroimage, 42(2), 936–944.

Glasser, M. F., Coalson, T. S., Robinson, E. C., Hacker, C. D., Harwell, J., Yacoub, E., Ugurbil, K., Andersson, J., Beckmann, C. F., Jenkinson, M., Smith, S. M., & Van Essen, D. C. (2016). A multimodal parcellation of human cerebral cortex. Nature, 536(7615), 171–178.

Gross, J., Baillet, S., Barnes, G. R., Henson, R. N., Hillebrand, A., Jensen, O., Jerbi, K., Litvak, V., Maess, B., & Oostenveld, R. (2013). Good practice for conducting and reporting MEG research. Neuroimage, 65, 349–363.

Hari, R. (1990). The neuromagnetic method in the study of the human auditory cortex. Advances in Audiology, 6, 222–282.

Hill, R. M., Boto, E., Holmes, N., Hartley, C., Seedat, Z. A., Leggett, J., Roberts, G., Shah, V., Tierney, T. M., & Woolrich, M. W. (2019). A tool for functional brain imaging with lifespan compliance. Nature Communications, 10(1), 1–11.

Hill, R. M., Boto, E., Rea, M., Holmes, N., Leggett, J., Coles, L. A., Papastavrou, M., Everton, S. K., Hunt, A. E., Sims, D., Osborne, J., Shah, V., Bowtell, R., & Brookes, M. J. (2020). Multi-channel whole-head OPM-MEG: Helmet design and a comparison with a conventional system. NeuroImage, 219, 116995.

Hillebrand, A., & Barnes, G. R. (2005). Beamformer analysis of MEG data. International Review of Neurobiology, 68, 149–171.

Holmes, N., Leggett, J., Boto, E., Roberts, G., Hill, R. M., Tierney, T. M., Shah, V., Barnes, G. R., Brookes, M. J., & Bowtell, R. (2018). A bi-planar coil system for nulling background magnetic fields in scalp mounted magnetoencephalography. NeuroImage, 181, 760–774.

Holmes, N., Tierney, T. M., Leggett, J., Boto, E., Mellor, S., Roberts, G., Hill, R. M., Shah, V., Barnes, G. R., Brookes, M. J., & Bowtell, R. (2019). Balanced, bi-planar magnetic field and field gradient coils for field compensation in wearable magnetoencephalography. Scientific Reports, 9(1), 14196.

Iivanainen, J., Stenroos, M., & Parkkonen, L. (2017). Measuring MEG closer to the brain: Performance of on-scalp sensor arrays. NeuroImage, 147, 542–553.

Iivanainen, J., Zetter, R., Grön, M., Hakkarainen, K., & Parkkonen, L. (2019). On-scalp MEG system utilizing an actively shielded array of optically-pumped magnetometers. NeuroImage, 194, 244–258.

Iivanainen, J., Zetter, R., & Parkkonen, L. (2020). Potential of on-scalp MEG: Robust detection of human visual gamma-band responses. Human Brain Mapping, 41(1), 150–161.

Jungnickel, E., & Gramann, K. (2016). Mobile Brain/Body Imaging (MoBI) of Physical Interaction with Dynamically Moving Objects. Frontiers in Human Neuroscience, 10.

Kessler, K., Seymour, R. A., & Rippon, G. (2016). Brain oscillations and connectivity in autism spectrum disorders (ASD): New approaches to methodology, measurement and modelling. Neuroscience & Biobehavioral Reviews, 71, 601–620.

Kowalczyk, A. U., Bezsudnova, Y., Jensen, O., & Barontini, G. (2021). Detection of human auditory evoked brain signals with a resilient nonlinear optically pumped magnetometer. NeuroImage, 226, 117497.

Leske, S., & Dalal, S. S. (2019). Reducing power line noise in EEG and MEG data via spectrum interpolation. Neuroimage, 189, 763–776.

Litvak, V., Eusebio, A., Jha, A., Oostenveld, R., Barnes, G. R., Penny, W. D., Zrinzo, L., Hariz, M. I., Limousin, P., Friston, K. J., & Brown, P. (2010). Optimized beamforming for simultaneous MEG and intracranial local field potential recordings in deep brain stimulation patients. NeuroImage, 50(4), 1578–1588.

Litvak, V., Mattout, J., Kiebel, S., Phillips, C., Henson, R., Kilner, J., Barnes, G., Oostenveld, R., Daunizeau, J., Flandin, G., & others. (2011). EEG and MEG data analysis in SPM8. Computational Intelligence and Neuroscience, 2011.

Maris, E., & Oostenveld, R. (2007). Nonparametric statistical testing of EEG- and MEG-data. Journal of Neuroscience Methods, 164(1), 177–190.

Medvedovsky, M., Taulu, S., Bikmullina, R., & Paetau, R. (2007). Artifact and head movement compensation in MEG. Neurol. Neurophysiol. Neurosci, 4(4).

Mellor, S., Tierney, T.M., O’Neill, G.C., Alexander, N. Seymour, R.A., … Barnes, G. (2021). Magnetic field mapping and correction for moving OP-MEG. BioRxiv.

Meyer, S. S., Bonaiuto, J., Lim, M., Rossiter, H., Waters, S., Bradbury, D., Bestmann, S., Brookes, M., Callaghan, M. F., & Weiskopf, N. (2017). Flexible head-casts for high spatial precision MEG. Journal of Neuroscience Methods, 276, 38–45.

Nardelli, N. V., Perry, A. R., Krzyzewski, S. P., & Knappe, S. A. (2020). A conformal array of microfabricated optically-pumped first-order gradiometers for magnetoencephalography. EPJ Quantum Technology, 7(1), 11.

Nichols, T. E., & Holmes, A. P. (2002). Nonparametric permutation tests for functional neuroimaging: A primer with examples. Human Brain Mapping, 15(1), 1–25.

Oostenveld, R., Fries, P., Maris, E., & Schoffelen, J.-M. (2011). FieldTrip: Open source software for advanced analysis of MEG, EEG, and invasive electrophysiological data. Computational Intelligence and Neuroscience, 2011, 1.

Osborne, J., Orton, J., Alem, O., & Shah, V. (2018). Fully integrated standalone zero field optically pumped magnetometer for biomagnetism. Steep Dispersion Engineering and Opto-Atomic Precision Metrology XI, 10548, 105481G.

Peirce, J. W. (2009). Generating stimuli for neuroscience using PsychoPy. Frontiers in Neuroinformatics, 2, 10.

Popov, T., Oostenveld, R., & Schoffelen, J. M. (2018). FieldTrip Made Easy: An Analysis Protocol for Group Analysis of the Auditory Steady State Brain Response in Time, Frequency, and Space. Frontiers in Neuroscience, 12.

Roberts, G., Holmes, N., Alexander, N., Boto, E., Leggett, J., Hill, R. M., Shah, V., Rea, M., Vaughan, R., Maguire, E. A., Kessler, K., Beebe, S., Fromhold, M., Barnes, G. R., Bowtell, R., & Brookes, M. J. (2019). Towards OPM-MEG in a virtual reality environment. NeuroImage, 199, 408–417.

Roberts, T. P. L., Cannon, K. M., Tavabi, K., Blaskey, L., Khan, S. Y., Monroe, J. F., Qasmieh, S., Levy, S. E., & Edgar, J. C. (2011). Auditory Magnetic Mismatch Field Latency: A Biomarker for Language Impairment in Autism. Biological Psychiatry, 70(3), 263–269.

Rojas, D. C., & Wilson, L. B. (2014). γ-band abnormalities as markers of autism spectrum disorders. Biomarkers in Medicine, 8(3), 353–368.

Schiza, E., Matsangidou, M., Neokleous, K., & Pattichis, C. S. (2019). Virtual Reality Applications for Neurological Disease: A Review. Frontiers in Robotics and AI, 6.

Schoffelen, J.-M., Hultén, A., Lam, N., Marquand, A. F., Uddén, J., & Hagoort, P. (2017). Frequency-specific directed interactions in the human brain network for language. Proceedings of the National Academy of Sciences, 114(30), 8083–8088.

Sekihara, K., Nagarajan, S. S., Poeppel, D., & Marantz, A. (2002). Performance of an MEG adaptive-beamformer technique in the presence of correlated neural activities: Effects on signal intensity and time-course estimates. IEEE Transactions on Biomedical Engineering, 49(12), 1534–1546.

Sekihara, Kensuke, Nagarajan, S. S., Poeppel, D., Marantz, A., & Miyashita, Y. (2002). Application of an MEG eigenspace beamformer to reconstructing spatio-temporal activities of neural sources. Human Brain Mapping, 15(4), 199–215.

Seymour, R. A., Rippon, G., Gooding-Williams, G., Sowman, P. F., & Kessler, K. (2020). Reduced auditory steady state responses in autism spectrum disorder. Molecular Autism, 11(1), 1–13.

Taulu, S., & Hari, R. (2009). Removal of magnetoencephalographic artifacts with temporal signal-space separation: Demonstration with single-trial auditory-evoked responses. Human Brain Mapping, 30(5), 1524–1534.

Tierney, T. M., Levy, A., Barry, D. N., Meyer, S. S., Shigihara, Y., Everatt, M., Mellor, S., Lopez, J. D., Bestmann, S., & Holmes, N. (2021a). Mouth magnetoencephalography: A unique perspective on the human hippocampus. NeuroImage, 225, 117443.

Tierney, T. M., Alexander, N., Mellor, S., Holmes, N., Seymour, R., O’Neill, G. C., Maguire, E. A., & Barnes, G. R. (2021b). Modelling optically pumped magnetometer interference in MEG as a spatially homogeneous magnetic field. BioRxiv.

Tierney, T. M., Holmes, N., Mellor, S., López, J. D., Roberts, G., Hill, R. M., Boto, E., Leggett, J., Shah, V., Brookes, M. J., Bowtell, R., & Barnes, G. R. (2019). Optically pumped magnetometers: From quantum origins to multi-channel magnetoencephalography. NeuroImage, 199, 598–608.

Tierney, T. M., Holmes, N., Meyer, S. S., Boto, E., Roberts, G., Leggett, J., Buck, S., Duque-Muñoz, L., Litvak, V., & Bestmann, S. (2018). Cognitive neuroscience using wearable magnetometer arrays: Non-invasive assessment of language function. NeuroImage, 181, 513–520.

Van Veen, B. D., van Drongelen, W., Yuchtman, M., & Suzuki, A. (1997). Localization of brain electrical activity via linearly constrained minimum variance spatial filtering. IEEE Transactions on Biomedical Engineering, 44(9), 867–880.

Vivekananda, U., Mellor, S., Tierney, T. M., Holmes, N., Boto, E., Leggett, J., Roberts, G., Hill, R. M., Litvak, V., & Brookes, M. J. (2020). Optically pumped magnetoencephalography in epilepsy. Annals of Clinical and Translational Neurology, 7(3), 397–401.

